# Effector prediction and characterization in the oomycete pathogen *Bremia lactucae* reveal host-recognized WY domain proteins that lack the canonical RXLR motif

**DOI:** 10.1101/679787

**Authors:** Kelsey Wood, Munir Nur, Juliana Gil, Kyle Fletcher, Kim Lakeman, Ayumi Gothberg, Tina Khuu, Jennifer Kopetzky, Archana Pandya, Mathieu Pel, Richard Michelmore

## Abstract

Pathogens infecting plants and animals use a diverse arsenal of effector proteins to suppress the host immune system and promote infection. Identification of effectors in pathogen genomes is foundational to understanding mechanisms of pathogenesis, for monitoring field pathogen populations, and for breeding disease resistance. We identified candidate effectors from the lettuce downy mildew pathogen, *Bremia lactucae*, using comparative genomics and bioinformatics to search for the WY domain. This conserved structural element is found in *Phytophthora* effectors and some other oomycete pathogens; it has been implicated in the immune-suppressing function of these effectors as well as their recognition by host resistance proteins. We predicted 54 WY domain containing proteins in isolate SF5 of *B. lactucae* that have substantial variation in both sequence and domain architecture. These candidate effectors exhibit several characteristics of pathogen effectors, including an N-terminal signal peptide, lineage specificity, and expression during infection. Unexpectedly, only a minority of *B. lactucae* WY effectors contain the canonical N-terminal RXLR motif, which is a conserved feature in the majority of cytoplasmic effectors reported in *Phytophthora* spp. Functional analysis effectors containing WY domains revealed eleven out of 21 that triggered necrosis, which is characteristic of the immune response on wild accessions and domesticated lettuce lines containing resistance genes. Only two of the eleven recognized effectors contained a canonical RXLR motif, suggesting that there has been an evolutionary divergence in sequence motifs between genera; this has major consequences for robust effector prediction in oomycete pathogens.

**Author Summary:** There is a microscopic battle that takes place at the molecular level during infection of plants and animals by pathogens. Some of the weapons that pathogens battle with are known as “effectors,” which are secreted proteins that enter host cells to alter physiology and suppress the immune system. Effectors can also be a liability for plant pathogens because plants have evolved ways to recognize these effectors, triggering a defense response leading to localized cell death, which prevents the spread of the pathogen. Here we used computer models to predict effectors from the genome of *Bremia lactucae*, the causal agent of lettuce downy mildew. Three effectors were demonstrated to suppress the basal immune system of lettuce. Eleven effectors were recognized by one or more resistant lines of lettuce. In addition to contributing to our understanding of the mechanisms of pathogenesis, this study of effectors is useful for breeding disease resistant lettuce, decreasing agricultural reliance on fungicides.

## Introduction

The phylum Oomycota includes some of the most devastating pathogens of both plants and animals [1]. Although oomycetes resemble fungi in their filamentous growth and infection structures, they are more closely related to brown algae than to fungi [2]. Notable oomycetes include the plant pathogens causing late blight of potato (*Phytophthora infestans*) [3], sudden oak death (*Phytophthora ramorum*) [4], root rot (*Phytophthora* [5] and *Pythium* spp. [6,7]), white blister rust of *Brassica* spp. (*Albugo* spp.) [8], and downy mildews (e.g. *Bremia, Peronospora, Plasmopara* spp.) [9], as well as several important animal pathogens infecting fish (*Saprolegnia* spp.), shellfish (*Aphanomyces astaci*), and mammals, including humans (*Pythium insidiosum*) [1,10]. Many types of plant and animal pathogens, including the oomycetes, secrete proteins known as effectors to promote virulence by manipulating the physiology of the host cells and by suppressing the host immune system [11]. In plants, effectors are also determinants of resistance or susceptibility through their direct or indirect interactions with cognate nucleotide-binding leucine rich repeat proteins (NLRs) encoded by resistance genes [12]. Effectors are secreted from the pathogen and may act extracellularly or they may be translocated into the cytoplasm [11]. One class of translocated effectors from plant pathogenic oomycetes of the class Peronosporales, which includes *Phytophthora* and the downy mildews, are the RXLR effectors, so called for their N-terminal motif usually consisting of arginine, followed by any amino acid, then followed by leucine and arginine. RXLR effectors also contain an N-terminal signal peptide, which designates them for extracellular transport by way of the endoplasmic reticulum and Golgi apparatus [13]. The RXLR motif is often associated with a downstream EER motif, both of which have been associated with secretion and/or translocation of effectors into the plant cell [13,14]. For some RXLR effectors, such as Avr3a, the RXLR motif has been shown to be cleaved just prior to the EER sequence and therefore plays a role in secretion rather than uptake into the host cell [15]. The RXLR motif is similar in sequence to the PEXEL motif (RXLX[EDQ]) of the distantly related malaria pathogen (*Plasmodium falciparum*) [16] and the TEXEL motif of *Toxoplasma gondii* (RRLXX) [17], both of which are required for proteolytic modification in the endoplasmic reticulum and destine effector proteins for specialized export out of the cell [18].

The downy mildews and the related *Phytophthora* species have different lifestyles (obligate biotrophy *vs.* facultative hemibiotrophy) [19]; however, their effectors share similar features. Many effectors in the Peronosporales have RXLR and EER motifs; although several alternative sequences to RXLR have been found in downy mildews, including RVRN (ATR5 from *Hyaloperonospora arabidopsidis*) [20], QXLR (*Pseudoperonospora cubensis*) [21], GKLR (*Bremia lactucae*) [22,23], and RXLK (*Plasmopara halstedii*) [24]. The C-terminal effector domains of RXLR effectors from *Phytophthora* and downy mildews also share some common sequence motifs and structural features, such as the 24 to 30 amino acid W, Y, and L motifs, which were first identified bioinformatically [25]. Structural analysis on four different RXLR effectors from *Phytophthora infestans* (Avr3a and PexRD2), *P. capsici* (Avr3a11), and the downy mildew pathogen *H. arabidopsidis* (ATR1) revealed that the W and Y motifs form an alpha-helical fold that may play a role in protein-protein interactions [26– 29]. This effector-associated fold, termed the WY domain after its conserved tryptophan and tyrosine residues, is structurally highly conserved between effectors from multiple Peronosporales species, despite sharing less than 20% sequence similarity across the whole domain [26][30]. The WY domain appears to be specific to the Peronosporales and was predicted to be present in nearly half of the RXLR effectors of *P. infestans* and a fourth of the RXLR effectors in *H. arabidopsidis* [26].

Functional studies of WY domain containing proteins have indicated that certain residues in the WY domain are essential for the immune suppressing functions of *P. sojae* Avr1b [31], *P. infestans* Avr3a [32], and *P. infestans* PexRD2 [33]. Furthermore, mutation of two conserved leucines in the WY domain of PexRD2 disrupted interaction with its target, MAPKKKε, consistent with this domain being important for protein-protein interactions [33]. WY domain containing proteins appear to interact with a variety of host targets, with Avr3a from *P. infestans* targeting the E3 ligase CMPG1 [32], *P. infestans* PexRD54 targeting potato autophagy-related protein ATG8 [34], PsAvh240 from *P. sojae* targeting an aspartic protease [35], and PSR2 from *P. sojae* and *P. infestans* suppressing host RNA silencing through interactions with dsRNA-binding protein DRB4 [36–38]. Mutation analysis of the regions encoding the seven individual WY domains of PSR2 demonstrated differential contributions of each domain to virulence of *P. sojae*, suggesting that the WY domain may act as a module during effector evolution [30]. In addition to its roles in immune suppression, the WY domain has been shown to be important for immune recognition of the effector by nucleotide binding-leucine rich repeat (NB-LRR) resistance proteins [39].

Effector annotation in oomycete genomes has often relied on sequence similarity to known effectors or on prediction of conserved motifs, such as the RXLR motif, or in the case of Crinklers, the LXLFLAK motif [3]. Due to the short length and degeneracy of the RXLR sequence, the motif occurs frequently by chance; therefore, there is a high false positive rate (>50%) using string-based searches [25]. HMM-based searches have much lower false positive rates, but the false negative rate may be higher if the genome of interest has diverged significantly from the species used to build the HMM. Downy mildews have a narrow host range and pathogenicity-related proteins are likely to show high lineage-specificity due to co-evolution with their hosts. There is already evidence that downy mildew effectors show divergence from the canonical RXLR motif [20,23,24], thus complementary approaches for effector prediction that utilize other conserved features, such as the WY domain, are necessary to fully characterize the repertoire. In addition, due to the importance of the WY domain in effector function [31–33], WY domain containing proteins may be better candidates for effectors than those containing only the RXLR motif.

To identify a more complete repertoire of candidate effectors in the reference genome of *B. lactucae* and to test whether the WY domain is informative for predicting effectors in the Peronosporales, we searched for this domain using an HMM built from sequences of the WY domain in three *Phytophthora* species [27]. This revealed additional effector candidates that had not been found using an RXLR-based search; similar results were also obtained for other downy mildews and well-studied *Phytophthora* species. These predicted WY proteins from *B. lactucae* had other signatures of oomycete effectors, such as presence of a secretion signal, N-terminal intrinsic disorder, lineage specificity, and expression during infection. A subset of the predicted WY effectors suppressed the host immune system, while others elicited programmed cell death in specific genotypes of lettuce, indicative of recognition by host resistance proteins. Therefore, searches for the WY domains are highly useful for identification of downy mildew effectors lacking the RXLR motif, which was previously considered canonical for effectors of the Peronosporales.

## Results

Our HMM search initially identified 59 candidate WY effectors encoded by the gene models predicted in the *B. lactucae* SF5 assembly [40]; however, three pairs of genes appeared to be allelic based on sequence similarity and read depth, leaving a total of 55 non-redundant genes encoding candidate effectors (Figure 1A; Supplemental Table 1). Signal peptides were predicted for 43 of these 55 proteins, of which two proteins had a predicted transmembrane helix outside of the signal peptide (Figure 1A). Several predicted WY proteins seemed to be missing their start codons due to N-terminal truncation when compared to close relatives. One of these predicted proteins, BLN06, was found to be missing a significant portion of the N-terminal sequence in SF5 compared to BL24, which was the source isolate for the cloning of *BLN06* reported in Pelgrom *et al*. [41].

**Figure 1.**
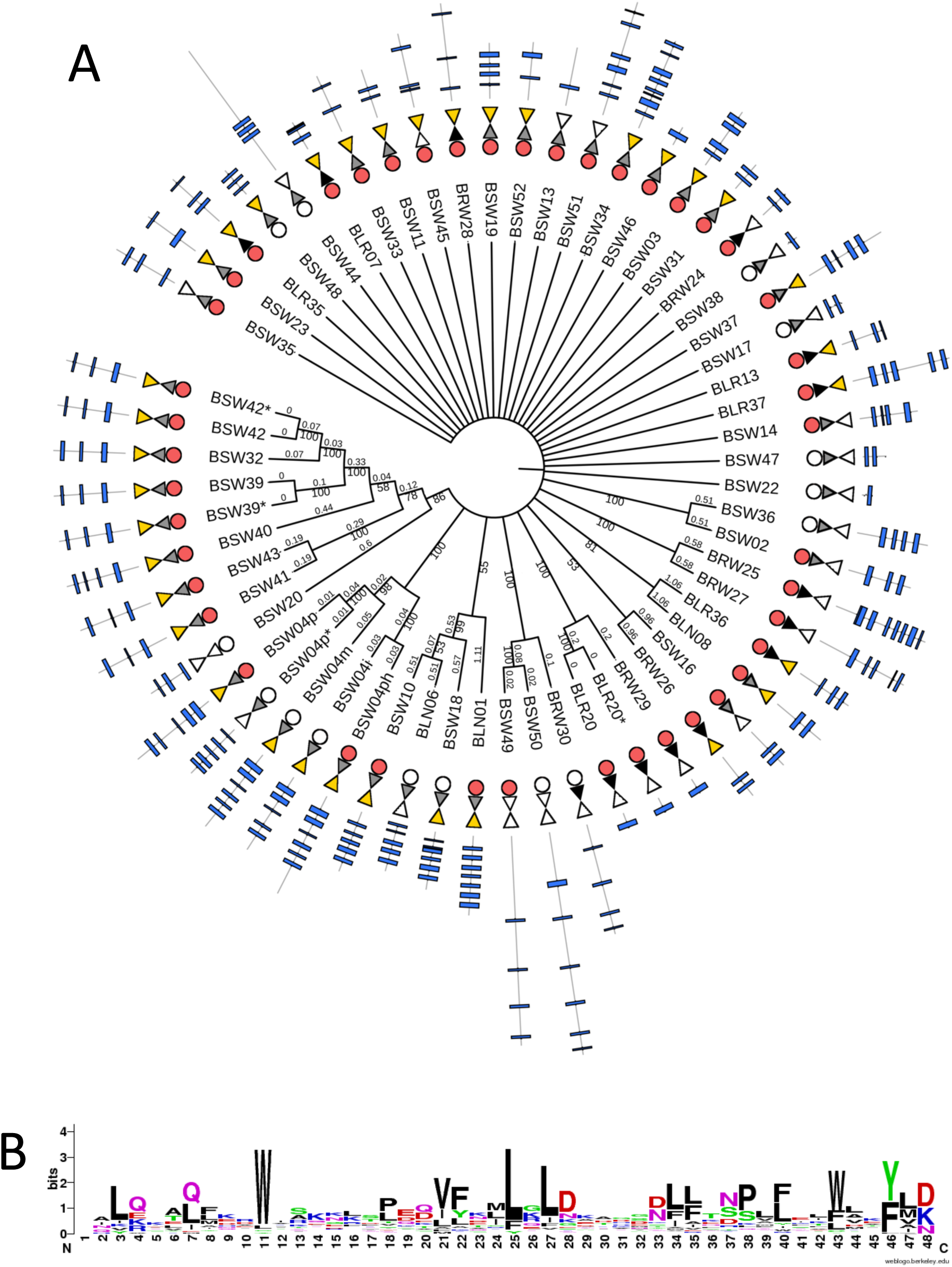
WY effector candidates from *B. lactucae* isolate SF5. (A) UPGMA consensus tree of the 59 predicted WY effectors from *B. lactucae* isolate SF5 based on whole protein amino acid sequence alignment using MUSCLE. Sequences that appear to be allelic are indicated with asterisks next to the sequence name. Bootstrap values and branch lengths are given for closely related proteins. Signal peptides are shown as circles (red circle for SP; white circle for no SP). RXLR motifs are shown as triangles (black triangle for RXLR ([RQGH]xLR or RXL[QKG]); grey triangle for degenerate RXLR ([RKHGQ][X]{0,1}[LMYFIV][RNK]), white triangle for no RXLR-like sequence) followed by an inverted triangle for EER motifs (yellow triangle for [DE][DE][RK], white triangle for no EER-like sequence). WY domain architecture is shown using blue rectangles, with a black line representing the total length of the protein and rectangle position representing the location of the WY motif as predicted by HMMer 3.0. (B) Sequence logo for the WY domain from *B. lactucae* built from a multiple sequence alignment of WY domains predicted by HMMer 3.0.

The N-terminal sequences of the 55 predicted WY effectors were examined for the RXLR motif using both HMMs and string searches. One protein was predicted to have an RXLR motif by the HMM and an additional 10 proteins were identified by a string search for [RQGH]xLR or RXL[QKG], while 33 proteins were predicted to have an EER motif using a string search for [DE][DE][EK] (Figure 1A). To find divergent RXLR-like motifs in the WY effector candidates, we searched for a highly degenerate pattern based on mutational studies of the RXLR motif [35] and natural RXLR variants reported for other downy mildews [5–9]. This revealed an additional 38 proteins with an RXLR-like motif within the first 60 amino acids after the signal peptide matching the pattern [RKHGQ][X]{0,1}[LMYFIV][RNK], many of which also had an EER motif (Figure 1A) (Supplemental Table 1). Therefore, the majority of candidate WY effectors in *B. lactucae* have a non-canonical RXLR motif. In the C-terminal domain, the WY effector candidates had one to seven WY domains per protein, with a diversity of domain architectures (Figure 1). The WY domain from *B. lactucae* showed high conservation of the characteristic conserved tryptophan (W) residue but appeared to show equal preference for tyrosine (Y) or phenylalanine (F) for the second characteristic residue of the domain (Fig. 1B). To investigate whether or not there are WY effector candidates that are lacking the canonical RXLR motif in other oomycetes, the predicted open reading frames (ORFs) from several published oomycete genomes were surveyed for the RXLR motif and the WY domain. Approximately half of the WY proteins predicted in seven downy mildew species lacked the RXLR motif; in six *Phytophthora* spp., the majority of predicted WY proteins had RXLR motifs, but between 9–21% of secreted WY proteins did not contain this motif (Fig. 2). Consequently, the repertoire of candidate WY effectors in other oomycetes may be heavily under-reported.

**Figure 2.**
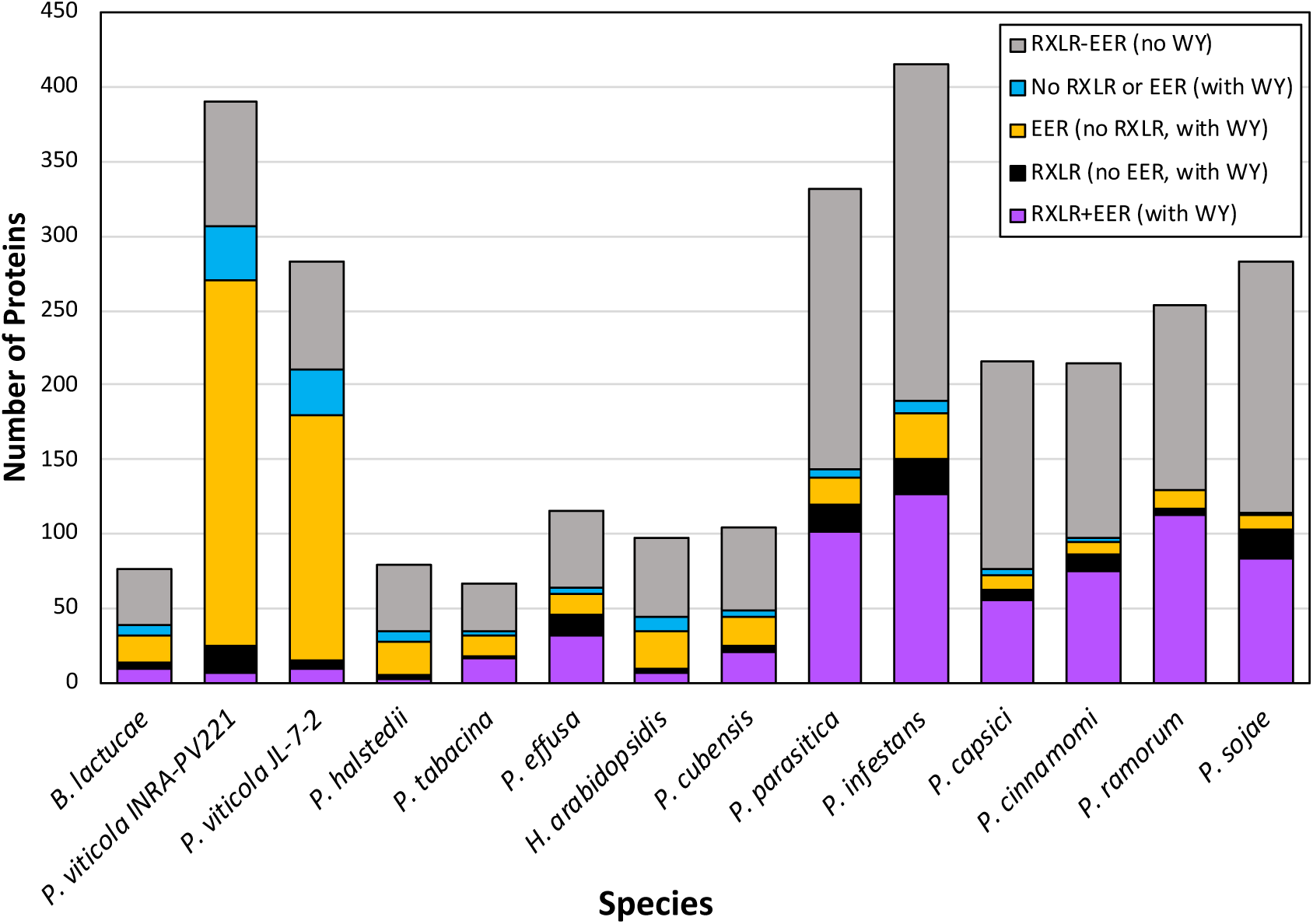
Distribution of predicted secreted WY effectors with or without RXLR and/or EER motifs in the genomes of downy mildew pathogens and *Phytophthora* species. HMMer was used to search predicted secretomes for each species for the WY domain. For the WY domaining containing proteins, the presence of RXLR and EER were determined by searching for [RQGH]XLR or RXL[QKG] within the first 60 amino acids after the signal peptide and [DE][DE][KR] within the first 100 amino acids after the signal peptide. For the non-WY domain containing proteins, the RXLR and EER was determined by searching for a strict “RXLR” with [DE][DE][KR], within the first 60 and 100 amino acids, respectively, plus proteins found by searching for the RXLR-EER domain using the RXLR-EER HMM from [14].

Intrinsic disorder had previously been reported to be a characteristic of the N-terminus of oomycete effectors containing the RXLR motif [42]. Therefore, we investigated whether the predicted degree of structural disorder was a characteristic of candidate WY effectors lacking a canonical RXLR. The predicted levels of intrinsic disorder were calculated for proteins containing RXLR motifs, for proteins containing WY domains but no RXLR motif, as well as for the entire predicted secretome for comparison. Proteins containing RXLR and/or WY domains had higher levels of intrinsic disorder at their N-termini after the highly ordered signal peptide than the entire set of secreted proteins (Fig. 3). Proteins containing a WY domain but lacking a canonical RXLR motif had on average more disordered N-termini than effectors that had RXLR but not a WY domain. The WY domain containing region had higher levels of predicted structure than the RXLR-containing proteins that lacked a WY domain and the secreted proteins as a whole (Fig. 3), consistent with the WY domains forming an α-helix bundled structure [27]. This predicted pattern of high intrinsic disorder at the N-terminus and high structure towards the C-terminus was consistently observed in all six downy mildews and six *Phytophthora* species analyzed (Suppl.Fig. 2). Therefore, a high level of intrinsic disorder is a consistent characteristic of the N-termini of oomycete effectors, regardless of whether they have a canonical RXLR motif; the functional significance of this remains to be investigated.

**Figure 3.**
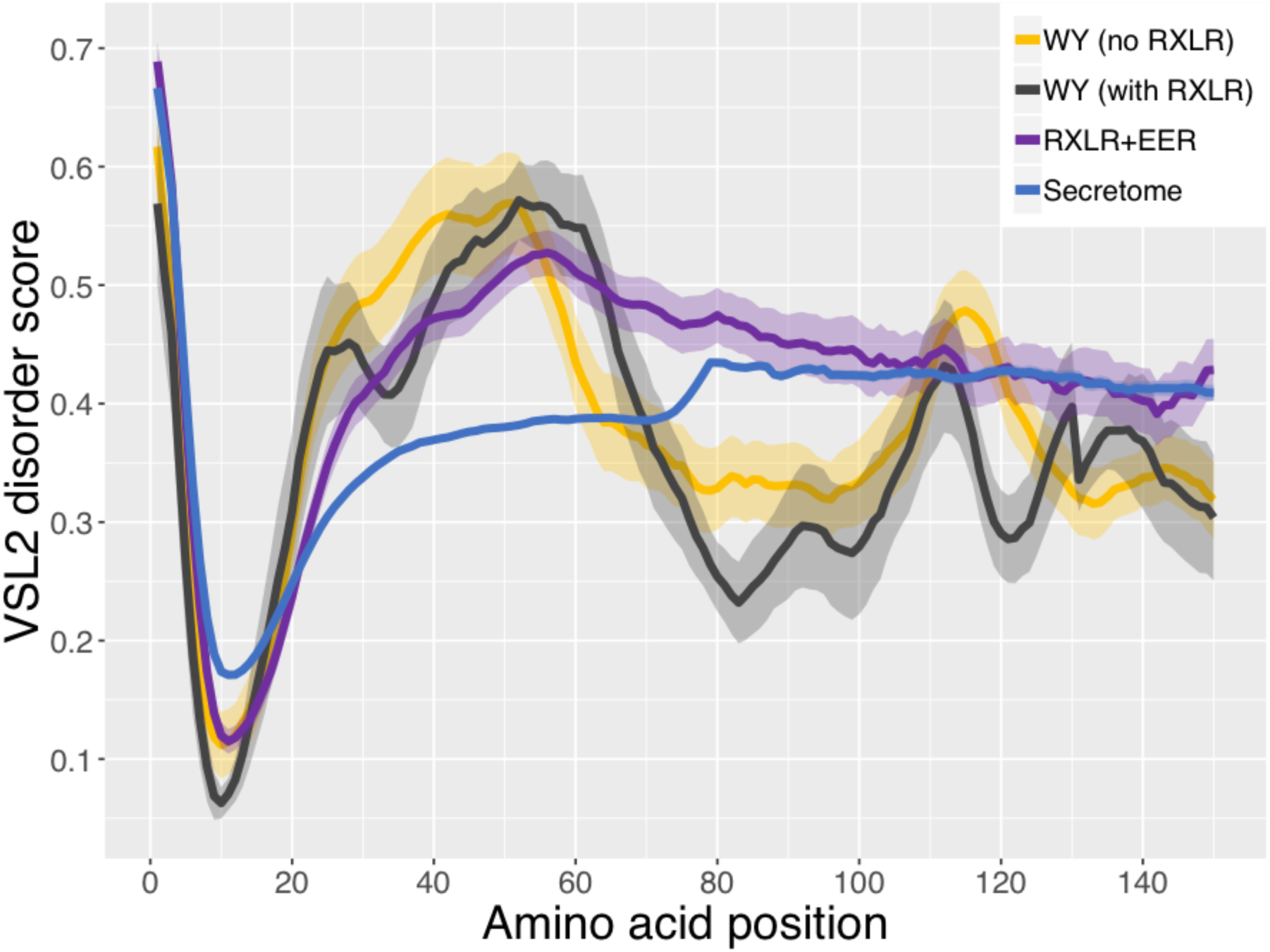
Intrinsic disorder in the first 150 N-terminal amino acid sequences of RXLR and/or WY containing candidate effectors in *B. lactucae*. Proteins were categorized as WY with no RXLR (yellow), WYs with RXLR (black), RXLR+EER (with or without WY, purple), and the total predicted secretome (blue). RXLR and EER motifs were as described in Fig 2. Average positional disorder scores [42] with standard error are shown for each group of proteins.

To evaluate lineage specificity of effectors due to co-evolution of pathogens with their hosts, we used BLAST to identify orthologs in other oomycete species. All of the 39 predicted secreted candidate WY effectors of *B. lactucae* had little sequence similarity to sequences in other genomes with the best BLASTP hit being only 46% identity with a protein from *P. infestans* (Fig. 4). Most of the proteins had best-hit identities between 20 to 30%, which is similar to the level of amino acid conservation between WY domains in different effectors [27].

**Figure 4.**
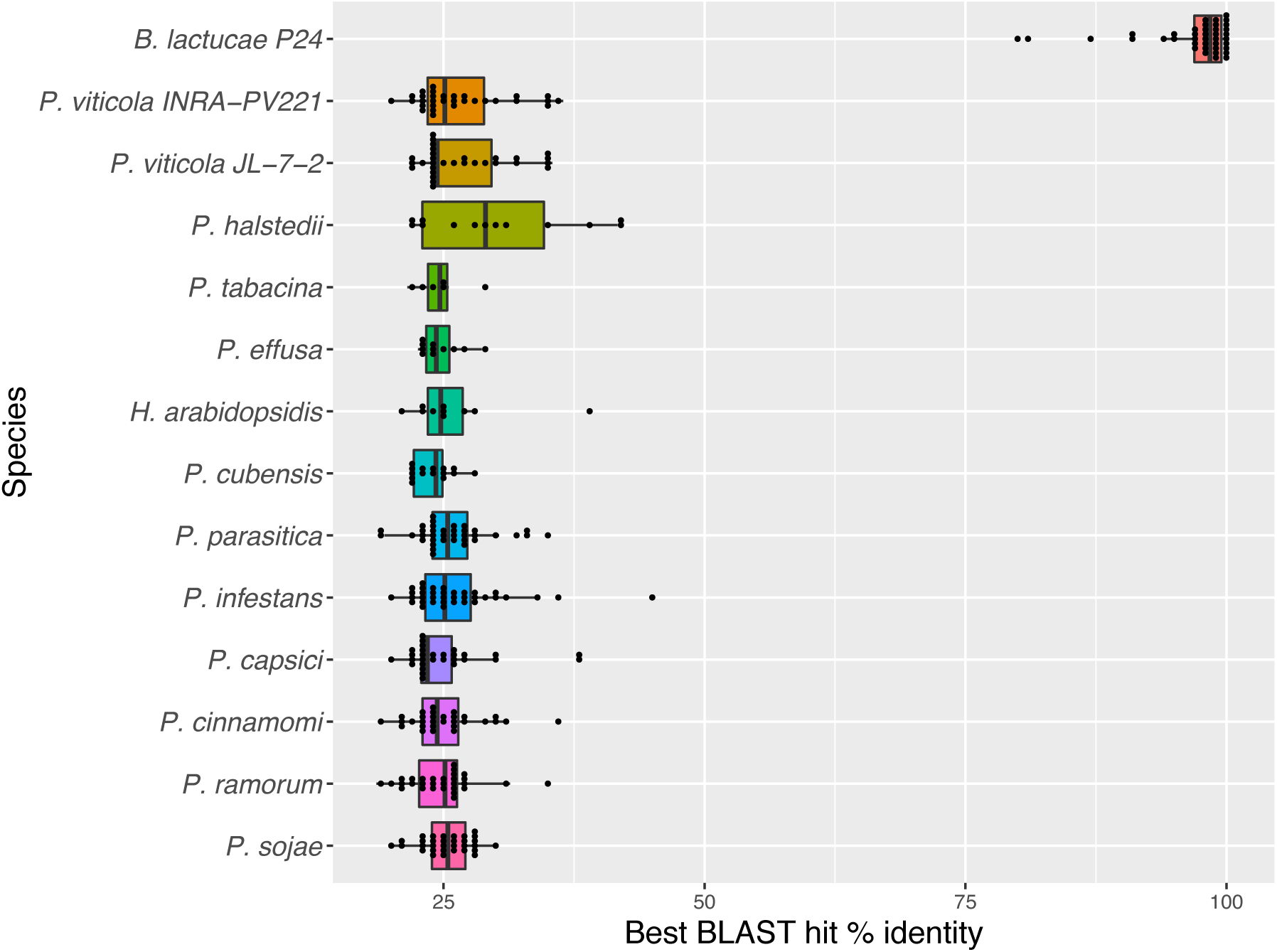
Lineage specificity of *B. lactucae* candidate secreted WY effectors. Secreted WY proteins predicted in isolate SF5 of *B. lactucae* were used as a query for BLASTp against other oomycete translated ORFs. Isolate C82P24 was queried for *B. lactucae.* The best BLAST hit percent identity for each protein was calculated. The box plot shows the distribution of the best BLAST hits per *B. lactucae* WY protein from each species, with each dot representing an individual data point. No hits were observed to *Albugo laibachii, Saprolegnia parasitica,* or *Pythium ultimum.*

A time-course RNA-seq experiment of lettuce seedlings infected with *B. lactucae* isolate SF5 was analyzed to investigate the expression of the WY effector candidates during infection. Expression was detected for all candidate WY encoding genes. The four mostly highly expressed WY containing effectors lacked a canonical RXLR motif (Fig. 5).

**Figure 5.**
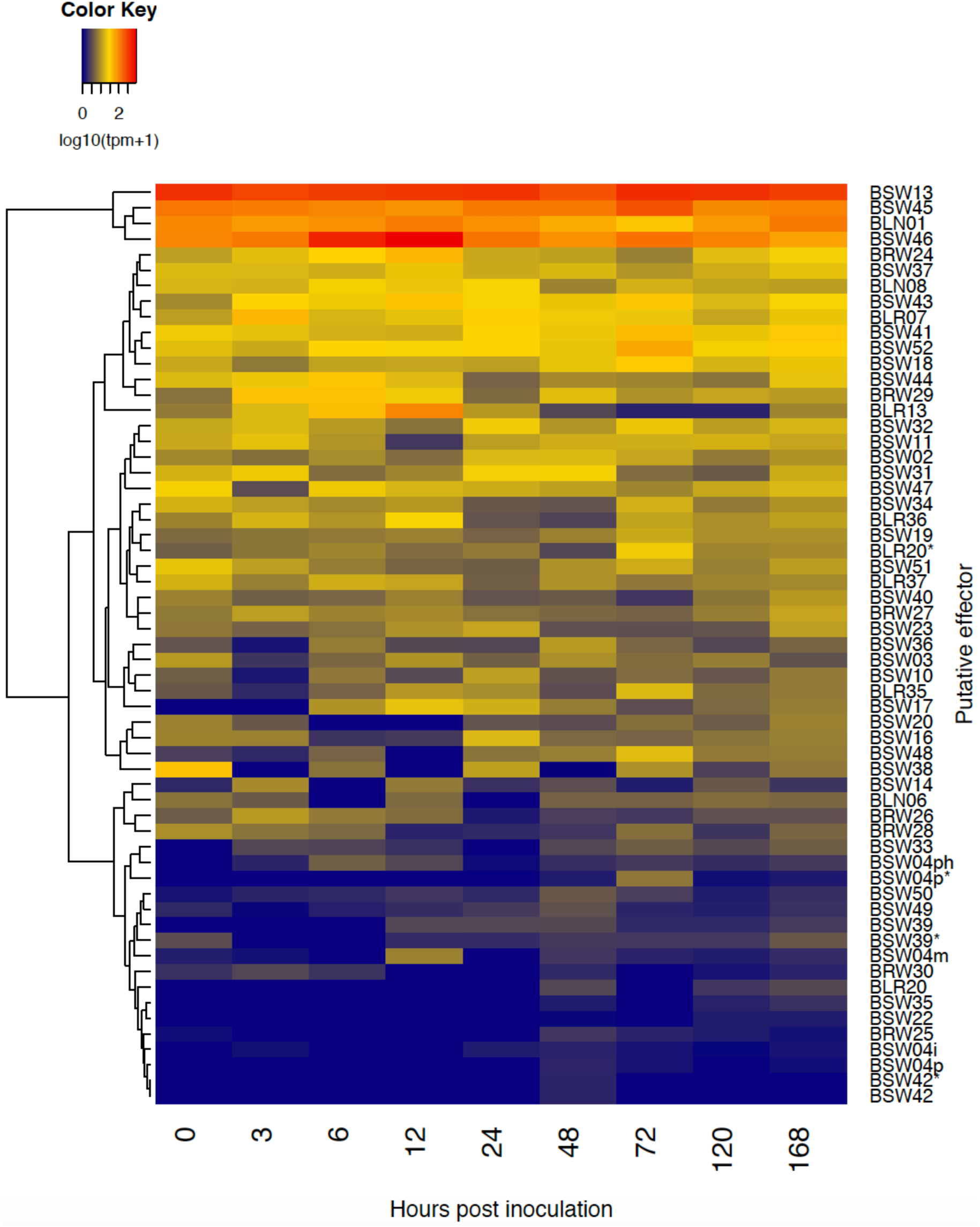
RNA-seq expression levels during infection for the *B. lactucae* WY effector candidates. The transcripts per million (TPM) of each WY-encoding gene was normalized by the total number of reads per kilobase assigned to *B. lactucae* from RNAseq analysis of infected cotyledons over a seven-day time-course. This value was log10(+1) transformed to enable visualization. The four highest expressed WY encoding genes do not have recognizable RxLR translocation associated motifs.

To test whether *B. lactucae* WY effectors were recognized by the host immune system, 21 randomly-selected WY effectors that differed in their number of WY domains were chosen to be screened against lettuce germplasm. Genes were cloned from amplified genomic DNA to attempt to capture both alleles of each effector from the heterozygous isolate SF5 [40]. In order to capture additional allelic diversity, some effectors were also cloned from the heterokaryotic isolate C82P24 [40]. Two alleles were obtained for many effectors; in addition, chimeric sequences appeared to be obtained for several genes. Clones of all unique sequences for each effector were retained because they could be informative for dissecting the sequence basis of host recognition. This resulted in multiple distinct clones (wildtype alleles plus chimeric sequences) of some effectors.

We screened 215 different accessions of wild and cultivated lettuce (Suppl. File 2) using *Agrobacterium*-mediated transient expression for the elicitation of cell death for their reactions to clones representing the 21 WY candidate effectors. These accessions collectively express the majority of the known *Dm* genes as well as new resistance factors [36]. Eleven of the 21 WY effectors were recognized by one or more accessions (Fig. 6). Alleles of the same effector showed similar reactions except for three genes (Fig. 6). The truncated version of BLN06 cloned from SF5 did not trigger cell death in LS102, NunDm17, or RYZ2146 (Suppl. Figure 3) in contrast to BLN06 cloned from BL24 [41], suggesting that the recognition of this effector by these genotypes may be determined by the N-terminal region of the effector after the secretion signal.

**Figure 6.**
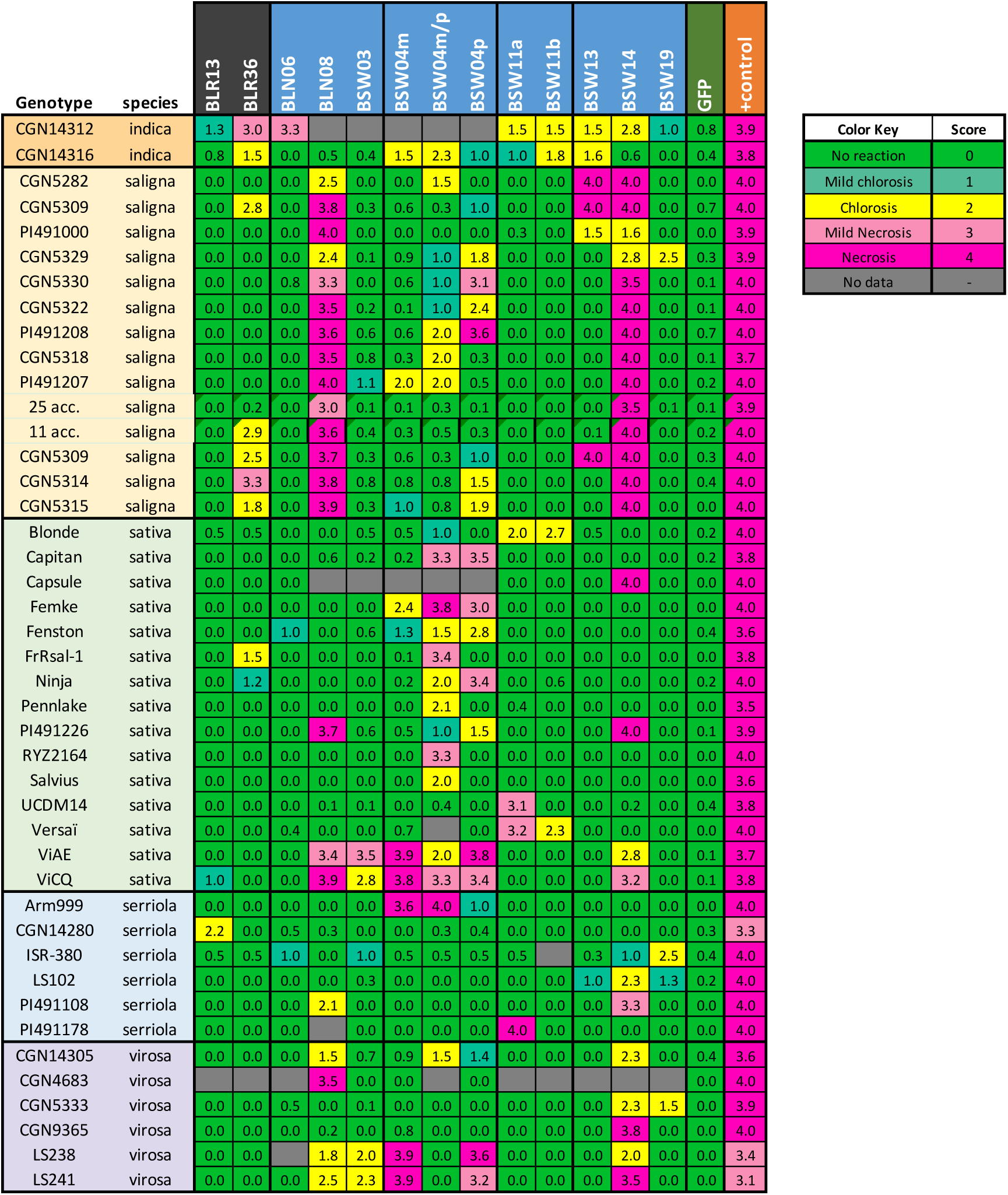
Results of the screen for recognition of *B. lactucae* WY effector by diverse genotypes of lettuce. Candidate effectors were expressed in lettuce leaves using Agroinfiltration and leaves were scored for their reaction four to seven days post-infiltration. Qualitative scores were converted into numeric scores for data analysis: 0 = no reaction (green), 1 = mild chlorosis (blue-green), 2 = chlorosis (yellow), 3 = mild necrosis or mixture of chlorosis and necrosis (pink), and 4 = full necrosis (magenta). The figure shows average scores for each effector on each genotype; only accessions and effectors that had a necrotic or chlorotic response are shown. Scores for all accessions screened and sample sizes can be found at http://bremia.ucdavis.edu/BIL/BIL_interaction.php. Two groups of 28 and 11 accessions of *L. saligna* that had identical reactions are shown together; the individual accessions making up each group are shown in Supplementary Table 2.

The lines ViAE and ViCQ, which have introgressions from the wild lettuce species *L. virosa* [43], recognized five effectors: BLN08, BSW03, BSW04m, BSW04p, and BSW14, as well as the chimeric sequence BSW04m/p (Fig. 6A). The *L. virosa* accessions that were the resistance donors for ViAE and ViCQ also recognized these five effectors, but not the chimeric sequence BSW04m/p (Fig. 6B). Many *L. sativa* cultivars and a few genotypes of *L. saligna* and *L. serriola* were observed to have necrosis or yellowing in response to the chimeric effector BSW04m/p, suggestive of a non-specific reaction to this unnatural protein. Some of the genotypes that recognized BSW04m/p also recognized BSW04p, but not the paralog BSW04m. Three of these genotypes, Capitan, Ninja, and Femke, share the resistance gene *Dm11*; therefore, we tested two additional cultivars, Fila and Mondian, which also contain *Dm11* for recognition of BSW04p. These varieties also recognized BSW04m/p and BSW04p, but not BSW04m. Due to recognition of BSW04p by multiple varieties that contain *Dm11*, BSW04p is a candidate for the protein encoded by the *Avr11* gene. BLN08, BSW03, BSW04m, and BSW14 are candidate avirulence proteins for which cognate *Dm* genes have yet to be identified.

Bioinformatic prediction of subcellular localization using NucPred [44] suggested that BSW04p had a C-terminal nuclear localization signal (score 0.79). To investigate whether BSW04p was nuclear localized, N-terminal yellow fluorescent protein (YFP) fusions were expressed in lettuce using *Agrobacterium*-mediated transient assays. We also made N-terminal YFP fusions of two other effectors, BLN08 and BSW03, which lacked predicted nuclear localization signals. BSW04p was localized to the nucleus as predicted, while BLN08 and BSW03 were localized to the cytoplasm and/or periplasm (Fig. 7). Therefore, predictions of subcellular localization were accurate for these three effectors and may indicate the cellular location of their targets during infection.

**Figure 7.**
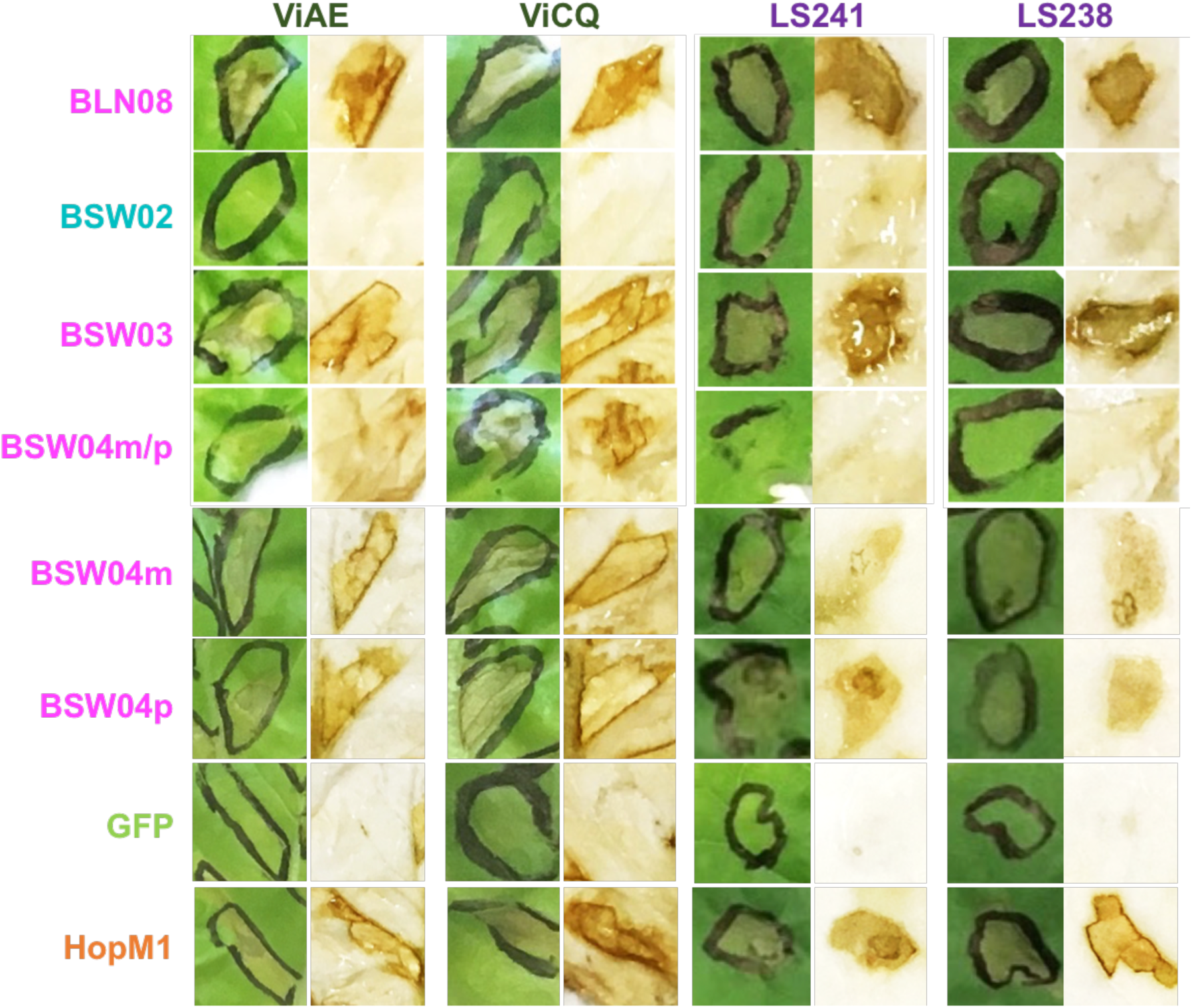
Example Agroinfiltration results of reactions of *L. sativa* ViAE and ViCQ and their progenitor R gene donors *L. virosa* LS238 and LS241 to several *B. lactucae* effectors. Photos are representative of a typical leaf. *B. lactucae* candidate effectors either elicited necrosis (brown areas) indicative of an immune recognition response (magenta text) or did not elicit a response (aqua text). GFP and HopM1 were used as negative and positive controls for necrosis, respectively. Leaves were collected five days post-infiltration and the first column for each accession shows the uncleared leaf tissue; the second column shows leaf tissue cleared in ethanol.

We also tested candidate effectors for their ability to suppress PAMP-triggered immunity (PTI). Twenty-one effectors were transiently expressed in *Nicotiana benthamiana* and the level of reactive oxygen species (ROS) production induced by flg22 was measured. Three effectors significantly suppressed ROS production to a similar extent as two known bacterial suppressors of PTI (Fig. 8). The level of induction of ROS by flg22 was significantly higher with some effectors; however, there was no induction of ROS in the absence of flg22. Therefore, in this assay, at least three effectors suppress PTI and some may actually increase PTI responsiveness.

**Figure 8.**
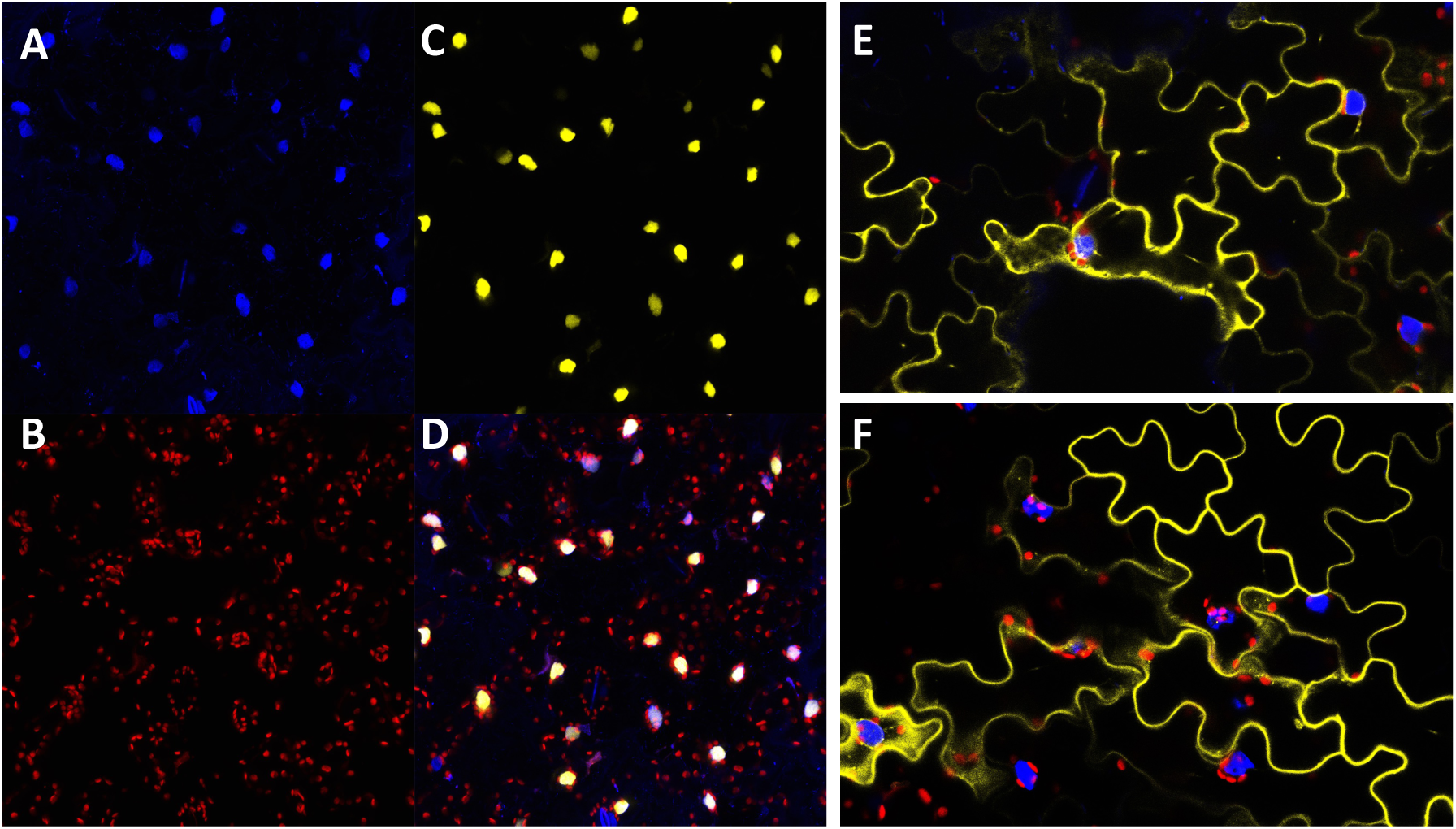
Subcellular localization of three *B. lactucae* WY effectors in lettuce. Confocal images of lettuce expressing YFP fusion proteins. For YFP-SW4, split images of (A) DAPI, (B) Chloroplast autofluorescence, (C) YFP, and (D) Merged show nuclear localization. For YFP-SW1 (E) and YFP-SW3 (F), merged images of DAPI (blue), chlorophyll autofluorescence (red), and YFP (yellow) show cytoplasmic or periplasmic localization.

**Figure 9.**
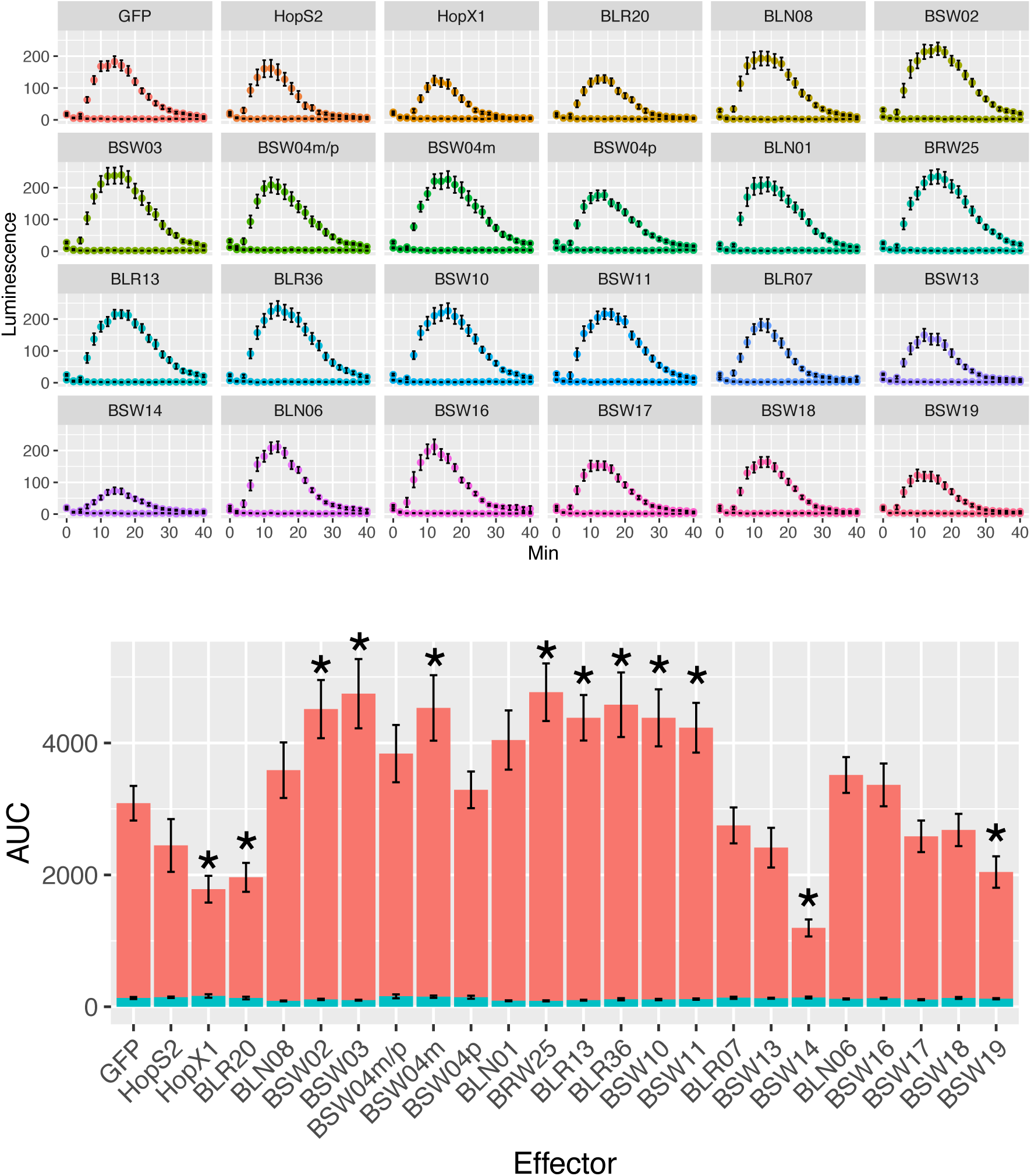
Effect of *B. lactucae* candidate WY effectors on flg22-triggered immune response. Luminescence plots measuring effect of candidate effectors on flg22-triggered PTI induction in *N. benthamiana* leaf discs. (A) Luminescence averaged across biological replicates (14–24 reps each) for each effector over time. (B) Bar plots of the average area under the curve (AUC) for luminescence over 40 minutes for each effector (N=14–24), with flg22 (red) or without flg22 (aqua). Asterisks indicate averages in the flg22+ condition that are statistically different from the average of GFP in the flg22+ condition (t-test, p < 0.05).

## Discussion

Effectors play a critical role in interactions between pathogens and their hosts. Accurate prediction and annotation of effector repertoires is foundational for functional genomics studies of pathogens. Due to their economic importance in agriculture, an increasing number of *Phytophthora* and downy mildew pathogens are being sequenced. Prior to this study, the majority of cloned avirulence genes have encoded proteins with a RXLR motif [45]. This was not surprising considering that most of these studies used the RXLR motif as their primary, or only, criterion to identify candidate effectors. Our results demonstrate that there are many candidate effectors containing the effector-related WY domain that lack the canonical RXLR motif, especially in the downy mildews, but also in *Phytophthora* spp. In *B. lactucae,* we have shown that these non-RXLR, WY domain containing effectors show characteristics of RXLR effectors, such as N-terminal intrinsic disorder, lineage specificity, and expression during infection. Furthermore, some of these proteins can act as suppressors of the host immune system, while others trigger the hypersensitive response in resistant host cultivars. Thus, despite lacking the canonical RXLR-motif, these WY domain containing proteins have both virulence and avirulence activity. Consequently, numerous candidate effector genes are likely to have been missed in the genomic analysis of other species within the Peronosporales. Bioinformatics pipelines for predicting effectors in both *Phytophthora* spp. and downy mildew pathogens should include an HMM for the WY domain because it may be more informative of function and result in fewer false positives than string searches for the RXLR motif.

Many WY effectors were initially predicted to not have a signal peptide and therefore not to be secreted; this could have been due to misannotation or reflect biological reality. Several had a predicted signal peptide downstream of the ATG in the gene model; these signal peptides started with a methionine, which was supported as the correct start in manual curation using RNAseq data. Therefore, studies that rely on the presence of a signal peptide when using predicted ORFs may be missing true effectors with misannotated start codons. Some WY effectors had clearly lost the signal peptide due to N-terminal deletion; this was observed by comparisons within effector families. The truncated effectors are unlikely to be functional because they are missing the secretion signal and would not therefore be secreted from the cell. Signal peptide loss may be an evolutionary strategy for the pathogen to evade recognition of effectors.

Although very few of the WY effectors from *B. lactucae* contained the canonical RXLR motif, nearly all of them contained a degenerate RXLR motif. However, the degenerate RXLR regular expression should not be used on its own to search for genes encoding RXLR proteins due to the high false positive rate (>50%). Functional studies are needed to ascertain which of these degenerate motifs can function similarly to the canonical RXLR motif in protein secretion [15]. Despite lacking the canonical RXLR motif, the WY effectors had other features similar to RXLR proteins such as high N-terminal intrinsic disorder and the presence an EER motif. N-terminal intrinsic disorder has been predicted for RXLR effectors of *Phytophthora* and is also a common feature of bacterial effectors [46]. The biological significance of these features remains to be studied, particularly whether these intrinsically disordered domains are important in post-translational modification or protein-protein interactions as is the case for intrinsically disordered regions in other organisms [47,48].

The number of WY domains per protein varied considerably, from one to seven in *B. lactucae*. Many pathogen effectors exhibit rapid evolution and divergence due to selective pressure of evolving host targets and host resistance proteins [25]. Duplication of domains may allow for evolution of novel effector functions or for evasion of host recognition while retaining function [30]. Variation in the number of repeated domains is reminiscent of NB-LRR proteins, which recognize effectors (and/or effector activity) and also have leucine-rich repeated domains [49,50]. Duplication of regions encoding WY domains may have happened within a gene through replication errors or between different genes through illegitimate recombination. The genomic sequences of the repeats will be analyzed in multiple isolates to reveal the origins of these duplications and domain expansion or contraction.

Analysis of the amino acid sequences of the WY domain in *B. lactucae* showed that it may be better considered as a W[Y/F] domain, due to the equal preference for phenylalanine and tyrosine for the second characteristic residue. This substitution has a BLOSUM score of 3, indicating that it is fairly common. Both tyrosine and phenylalanine are aromatic amino acids; however, the hydroxyl group on tyrosine makes it slightly bulkier, more polar, and introduces a potential phosphorylation site. Variation in this region is not uncommon in other oomycete species: the fifth of the seven domains in Psr2 of *P. sojae* has an F instead of a Y and is similar in structure to the single domain of ATR1 in *H. arabidopsidis* that has a cysteine at the Y position [30]. Further structural characterization is needed to reveal whether these substitutions alter protein structure and their biological function.

In oomycetes, WY domain containing effectors have been shown to have several functions including PTI suppression [51]. At least three WY effectors from *B. lactucae* were able to suppress the host immune system by interfering with pathogen-triggered production of ROS. Suppression of host defenses is critical to the survival of *B. lactucae* and therefore it is not surprising that multiple effectors target the basal immune system. Identification of the host targets of these effectors will determine which steps in the signal transduction cascade are modulated by each effector or may reveal candidates for susceptibility genes in the host that are required for successful proliferation of *B. lactucae*.

Effectors are powerful tools for the discovery and characterization of host resistance genes [52]. Eleven of the *B. lactucae* effectors tested were recognized by one or more lettuce lines. Their cognate R genes will be identified using mapping of segregating F_2:3_ and recombinant inbred line populations. Recognition of four *B. lactucae* effectors (BLG01, BLN08, BLR31, and BLR38) has been successfully mapped [23,41,53]. BLG01 and BLN08 have been shown to be recognized broadly by *L. saligna* [23,53]. Our study confirms the results for BLNO8 and revealed BSW14 as an additional effector recognized by *L. saligna*. BLN08 and BSW14 share little sequence similarity (besides containing WY domains). Non-host resistance in *L. saligna* is clearly complex [54], but these results show that it is mediated in part by recognition of multiple effectors.

Not all RXLR candidate effectors have WY domains and the presence of a WY domain is not required for the avirulence activity of all effectors; for example, the *B. lactucae* effectors BLG01 and BLG03 do not contain WY domains, yet they elicit an immune response in lettuce [23]. Structural elucidation of an RXLR effector lacking the WY motif, *H. arabidopsidis* ATR13, revealed that it contained a helical fold that was distinct from the WY fold [55]. It would be informative to determine and compare the protein structures of additional RXLR effectors to determine whether there are other conserved C-terminal domains that may be involved in effector function in a similar way as the WY domain.

Both RXLR and WY effectors provide tools for monitoring pathogen populations and effector-driven resistance breeding. Analysis of diverse, global isolates will allow the characterization of individual effector repertoires as well as the development of the pan-repertoire for a whole pathogen species. Effectors also will be highly instrumental in cloning their cognate resistance genes as well as the identification of effector targets in the host. In addition, screens for resistance using transient expression of individual effectors will allow the pyramiding of resistance genes with different specificities that will maximize the evolutionary hurdle for the pathogen to become virulent. Ultimately, knowledge of effector repertoires will allow data-driven deployment of resistance genes leading to more durable disease resistance [56].

## Materials and Methods

### Effector prediction

To search for the WY domain, a Hidden Markov Model (HMM) was built using HMMer v3.1 [57] based on the inferred amino acid sequences of WY domains from *P. infestans, P. soja*e, and *P. ramorum.* These sequences were obtained from the supplemental material of Boutemy *et al*. [27] who predicted genes encoding WY domains based on motif searches of candidate RXLR effectors and identified a 49 amino acid long motif that spanned the WY domain in the crystal structures of Avr3a11 and PexRD2 [26]. Candidate WY effectors were predicted using this HMM to search translated predicted ORF sequences (>80 amino acids) as well as gene models in the draft genome of *B. lactucae* isolate SF5 [40]. Sequences with a positive HMM bit score were considered to be putative WY domain effectors as in [27]. Signal peptide prediction was performed on candidate WY effectors using SignalP v 4.1 [58] and PhobiusSP [59]. Default settings were altered for SignalP v 4.1 to have sensitivity similar to SignalP v 3.0. Output was compared between SignalP v 4.1 sensitive and the combination of SignalP4.0 + SignalP 3.0, and identical results were obtained from the two methods. Gene models were more accurate for predicting signal peptides than translated ORFs — in part due to misannotated start codons upstream of the probable true start codon and signal peptides in the ORFs. However, on several occasions the gene model was missing a signal peptide found in the ORF – these gene models were manually updated to reflect this likely true start site. SignalP v 4.1 in sensitive mode was better able to predict signal peptides in proteins with misannotated start codons compared to SignalP v 4.0 or v 5.0.

### RXLR prediction

A combination of string searches and HMMs were used to search for the RXLR motif in oomycete predicted WY proteins. The following strings were used based on variants observed in downy mildews: [RQGH]xLR or RXL[QKG] for RXLR and [DE][DE][KR] for EER. The Whisson *et al*. HMM [14] was also tested, although this did not reveal any more RXLR effector candidates. To search for additional non-canonical RXLR motifs in WY candidates, a highly degenerate string of [RKHGQ][X]{0,1}[LMYFIV][RNK] was used.

### Estimation of false positive rates for effector prediction

We determined false positive rates for each motif by analyzing multiple permutations of the non-redundant secretome using the same pipeline as described above for effector prediction. At least ten random permutations of the sequence space were created using the MEME fasta-shuffle-letters program (with a kmer size of 1) using peptides starting after the cleavage site identified by SignalP v4.1. The false positive rate for each motif was estimated as the average frequency of detection in the permutated sequences divided by the observed frequency in the original sequences. The estimated false positive rate for the RXLR and WY motif searches are given in Table 1.

**Table 1.** False positive rates for RXLR and WY HMM and string searches across 14 oomycete genomes

| Motif | String/HMM | False Positive Rate (Average +SD) | Range |
| --- | --- | --- | --- |
| RXLR | RXLR | 45.2% ± 3.4% | 32-54% |
| RXLR | [RQGH]XLR or RXL[QKG] | 56% ± 2% | 50-64% |
| RXLR | Whisson HMM [14] | 1.5% ± 0.8% | 0.1-3.3% |
| EER | [DE][DE][KR] | 49.4% ± 2.6% | 41-62% |
| RXLR-EER | RXLR + [DE][DE][KR] | 9.4% ± 3.0% | 4-16% |
| Degenerate RXLR | [RKHGQ][X]{0,1}[LMYFIV][RNK] | 74% ± 0.8% | 71-77% |
| WY | Boutemy HMM [27] | 0.05% ± 0.09% | 0-0.3% |

### Intrinsic Disorder

PONDR VSL2 [60] was used to calculate levels of intrinsic disorder for groups of candidate effectors, grouped by the presence of an RXLR-like motif, EER-like motif, and/or WY domain. PONDR VSL2 is a meta-predictor of protein disorder that utilizes neural networks and amino acid context to give a weighted disorder score at each amino acid position. The average sequence disorder was calculated at each amino acid position in each group of effectors and aligned on the first amino acid. Plots were generated from the average positional disorder scores calculated from all peptides in a group. The entire non-redundant, predicted secretome was used as a reference for comparison.

### Sequence comparison and determination of lineage specificity

To generate a neighbor-joining tree of the WY effector candidates, whole protein amino acid alignments were performed using MUSCLE 3.8.425 [61] implemented in Geneious 11.0.5 (http://www.geneious.com) with default settings. A UPGMA tree was built in Geneious with bootstrap resampling (100 replicates). The resulting tree was annotated using the Interactive Tree of Life [62] and stylistically refined in the GNU Image Manipulation Program (http://www.gimp.org). To build a sequence logo for the WY domain from *B. lactucae,* a multiple sequence alignment of WY domains was built using MUSCLE 3.8.425 [61], was manually corrected for alignment errors, and the sequence logo generated using Weblogo (http://weblogo.berkeley.edu/logo.cgi).

To determine if the candidate WY proteins are unique to *B. lactucae,* BLASTp [63] was used to search for orthologs in other oomycete species. BLASTp-based sequence comparisons with an e-value threshold of 0.01 were performed against the following oomycete genomes: *Albugo laibachii* [8], *H. arabidopsidis* [64], *P. capsici* [65], *P. infestans* [3], *Phytophthora parasitica* [66], *P. ramorum* [5], *P. sojae* [5], *Pythium ultimum* [7], *Saprolegnia parasitica* [67], *Pseudoperonospora cubensis* [21], *Plasmopara halstedii* [24], *Plasmopara viticola* JL-7-2 [68], INRA-PV221 [69], *Peronospora tabacina* [70], and *Peronospora effusa* [71].

### RNA-seq analysis

Messenger RNA was isolated from cotyledons of lettuce cv. Cobham Green infected with isolate SF5 of *B. lactucae* with Dynabeads™ mRNA DIRECT™ Purification kit (Thermo Fisher Scientific, Waltham, MA) per manufacturer recommendations for plant tissue. Library construction was done following the protocol of Zhong *et al*. [72]. The resulting libraries were sequenced in single-end mode in a HiSeq 3000 at the UC Davis DNA Technologies Core (http://dnatech.genomecenter.ucdavis.edu/). The quality of the libraries was assessed using FastQC V0.11.2 [73]. Bacterial and human contaminants were filtered with BWA-MEM [74] mapping against custom references of microbial and human databases. The remaining reads were mapped to a joint reference made up of the *L. sativa* cv. Salinas (GenBank: GCF_002870075.1) and *B. lactucae* isolate SF5 (GenBank: GCA_004359215.1) reference assemblies using STAR v2.6.0 [75]. STAR was run with the options “--sjdbOverhang 99 --sjdbGTFtagExonParentGene Parent --quantMode GeneCounts”. The strand specific read counts for each replicate were calculated from the reads per gene table output by STAR. Reads counts were normalized for gene length by dividing the read counts by the length of the gene it was mapped to (Reads Per Kilobase: RPK). The total RPK of the *B. lactucae* portion of each replicate was calculated and used to divide the RPK to calculate the transcripts per million (TPM) for each gene. A subset of the genes that encode putative WY domains were taken from this total, the average TPM for three replicates of each time point was calculated (excluding replicate 2 at 72 hours, due to low coverage) and the log10(1+average TPM) was calculated for each time point. These values were plotted using Heatmap2 [76] from the package gplots, clustering of putative effectors by expression was performed with hclust [77].

### Gateway cloning

Effector candidates without their signal peptide were cloned into the pEarleyGate100 (pEG100) vector for plant expression [78] using Gateway cloning (Thermo Fisher Scientific). Candidate effector genes were amplified from genomic DNA isolated from spores of *B. lactucae* isolate SF5 or C82P24 using Phusion High Fidelity Polymerase (Thermo Fisher Scientific) and primers (Suppl. File 3) that amplified each ORF after the predicted signal peptide cleavage site. The Kozak sequence (ACCATG) was added to the forward primer for correct translational initiation. PCR products were purified using polyethylene glycol (PEG) precipitation to remove primers and primer-dimers, recombined into pDONR207 using BP Clonase II (Thermo Fisher Scientific), and transformed into chemically competent *Escherichia coli* DH10B cells. The resulting entry clones were sequenced using primers designed to pDONR207 to confirm gene identity and identify alleles. The entry clones were then recombined into pEG100 using LR Clonase II and transformed into *E. coli* to generate expression clones. The expression clones were then transformed into *Agrobacterium tumefaciens* strain C58rif^+^ using electroporation. Kanamycin-resistant colonies were confirmed for the transgene using gene-specific primers.

### Agrobacterium-mediated transient assays

Agroinfiltration and transient expression experiments were performed using conditions optimized for lettuce [79]. *A. tumefaciens* was grown overnight from glycerol stocks and resuspended in 10 mM MgCl_2_ to OD=0.3–0.5. The youngest fully expanded leaves of 3 to 4-week-old greenhouse grown lettuce plants were infiltrated with the *A. tumefaciens* cultures using a needless syringe. Leaves were examined four to five days post-infiltration for signs of macroscopic cell death, indicative of immune recognition by host resistance proteins. *A. tumefaciens* containing pEG100 empty vector or pEG100:GFP were used as negative controls; *A. tumefaciens* containing T-DNAs that expressed HopM1 or AvrPto, which are *Pseudomonas syringae* effectors known to elicit cell death in lettuce [80], were used as positive controls. To control for false negatives, leaves that did not show cell death in response to the positive control HopM1 were excluded from the analysis. To control for false positives, leaves that showed cell death in response to the empty vector or GFP control were also excluded from the analysis. In addition, any cell death observed in the initial screening was confirmed by repeating infiltrations on at least four plants for effectors found to cause cell death.

### Subcellular localization prediction and characterization

Nuclear localization was predicted using NucPred [44] and nuclear localization motifs predicted using LOCALIZER [81]. For localization experiments, effectors were cloned without their signal peptide into pEG104 [31] as described above resulting in an N-terminal YFP fusion. Three to five days post-Agroinfiltration, lettuce leaves expressing N-terminal YFP fusions of effectors were examined for subcellular localization. Nuclear staining was performed by incubating cut leaf tissue in 18 nM DAPI in water for at least five minutes. Imaging was performed on a Zeiss LSM 710 laser scanning confocal microscope with a 40x objective lens.

### PTI suppression assay

Effectors were expressed in five-week-old plants of *N. benthamiana* using Agroinfiltration as described above. Two days post-infiltration, two leaf discs (3.8 mm) were taken from each infiltration site away from the leaf veins using a cork borer. Leaf discs were floated abaxial side up in 200 μL of distilled water in a 96-well white assay plate and incubated at room temperature for 24 hrs. To measure suppression of ROS production [82], the water was removed and 100 μL of assay solution was added to the leaf discs, which contained 17 μg/mL luminol (Sigma-Aldrich, St Louis, MO) and 10 μg/mL horseradish peroxidase type 6A (Sigma-Aldrich). One of the two paired leaf discs was exposed to 100 nM of flg22 peptide (flg+) and the other was not exposed to any elicitor as a control for endogenous ROS production. The known bacterial suppressors of PTI, HopS2, and HopX1 [83], were used as positive controls for suppression of ROS production; GFP was used as the negative control. Luminescence was measured on a FilterMax F5 Plate Reader (Molecular Devices, San Jose, CA) promptly after adding in the assay solution and was measured every two minutes for a total of 42 minutes. Each assay was replicated sixteen times.

## Acknowledgements

We thank Keri Cavanaugh, Alyssa Schweickert, Dasan Gann, Catherine Lopez, Amber Robbins, Natalie Hamada, Pauline Sanders, and Sebastian Reyes Chin Wo (all UC Davis) for wet lab and computational assistance. We thank Gitta Coaker (UC Davis) for assistance with the ROS assay. We thank Brigitte Maisonneuve (INRA, Montfavet, France) for providing the FrRsal-1, ViAE, and ViCQ lines of lettuce and Ales Lebeda (Palacký University, Olomouc, Czech Republic) and Alex Beharav (The Hebrew University, Tel Aviv, Israel) for lines of *L. saligna*. We thank Lien Bertier, Dasan Gann, Anne Giesbers, and Elizabeth Georgian for helpful comments on earlier versions of this manuscript. The work was supported by an NSF Graduate Fellowship and a USDA Fellowship #2018-67011-28053 to KW and the NSF/USDA AFRI Microbial Sequencing Program award #2009-65109-05925 to RWM.

## Author Contributions

KW, JG, MP, and RWM conceived and designed the experiments. KW, JG, TK, MN, AP, AG, JK, and KL performed the experiments. KW, JG, KF, TK, MN, AP, AG, JK, and KL analyzed the data. KW, JG, and RWM wrote the paper. All authors approved the final submission.

## Data sources

The reference genome assembly of *B. lactucae* is available from NCBI, GenBank ID: GCA_004359215.1. RNAseq reads of lettuce cv. Cobham Green infected with *B. lactucae* isolate SF5 is available at NCBI SRA BioProject PRJNA523226. Sequences of effector proteins are available in Supplementary data. The all data for interactions between effectors and individual lines of *Lactuca* spp. are available at http://bremia.ucdavis.edu/BIL/BIL_interaction.php.

## List of Supplemental Materials

**Supplemental Table 1.** RXLR-like sequences observed within 100 amino acids of the N-terminus of *B. lactucae* WY proteins, sorted by first amino acid.

**Supplemental Figure 1.** The number of RXLR-like motifs detected in the first 100 amino acids of the N-terminus of *B. lactucae* WY effectors.

**Supplemental Figure 2.** Intrinsic disorder of first 150 amino acids of RXLR and WY containing candidate effectors mined from other oomycete genomes.

**Supplemental Figure 3.** Agroinfiltration results from BLN06-SF5 and sequence comparison between BLN06 between SF5 and BL24.

**Supplemental File 1.** Sequences and NCBI reference numbers of the 59 WY proteins predicted from the *B. lactucae* SF5 genome assembly.

**Supplemental File 2.** Lettuce genotypes tested for recognition of *B. lactucae* effectors.

**Supplemental File 3.** Primers used in Gateway Cloning of the *B. lactucae* effectors.

## References

1. Derevnina L, Petre B, Kellner R, Dagdas YF, Sarowar MN, Giannakopoulou A, et al. Emerging oomycete threats to plants and animals. Philos Trans R Soc B Biol Sci. 2016;371. doi:10.1098/rstb.2015.0459

2. Baldauf SL. The Deep Roots of Eukaryotes. 2003;300: 1703–1707. doi:10.1126/science.1085544

3. Haas BJ, Kamoun S, Zody MC, Jiang RHY, Handsaker RE, Cano LM, et al. Genome sequence and analysis of the Irish potato famine pathogen Phytophthora infestans. Nature. 2009;461: 393–398. doi:10.1038/nature08358

4. Rizzo DM, Garbelotto M, Hansen EM. Phytophthora ramorum: Integrative Research and Management of an Emerging Pathogen in California and Oregon Forests. Annu Rev Phytopathol. 2005;43: 309–335. doi:10.1146/annurev.phyto.42.040803.140418

5. Tyler BM, Tripathy S, Zhang X, Dehal P, Jiang RHY, Aerts A, et al. Phytophthora genome sequences uncover evolutionary origins and mechanisms of pathogenesis. Science. 2006;313: 1261–1266. doi:10.1126/science.1128796

6. Adhikari BN, Hamilton JP, Zerillo MM, Tisserat N, Lévesque CA, Buell CR. Comparative genomics reveals insight into virulence strategies of plant pathogenic oomycetes. PLoS One. 2013;8: e75072. doi:10.1371/journal.pone.0075072

7. Lévesque CA, Brouwer H, Cano L, Hamilton JP, Holt C, Huitema E, et al. Genome sequence of the necrotrophic plant pathogen Pythium ultimum reveals original pathogenicity mechanisms and effector repertoire. Genome Biol. 2010;11: 1–22. doi:10.1186/gb-2010-11-7-r73

8. Kemen E, Gardiner A, Schultz-Larsen T, Kemen AC, Balmuth AL, Robert-Seilaniantz A, et al. Gene gain and loss during evolution of obligate parasitism in the white rust pathogen of Arabidopsis thaliana. PLoS Biol. 2011;9: e1001094. doi:10.1371/journal.pbio.1001094

9. Thines M, Choi Y. Evolution, Diversity, and Taxonomy of the Peronosporaceae, with Focus on the Genus Peronospora. Phytopathology. 2015;106: 6–18.

10. Phillips AJ, Anderson VL, Robertson EJ, Secombes CJ, van West P. New insights into animal pathogenic oomycetes. Trends Microbiol. 2007;16: 13–19. doi:10.1016/j.tim.2007.10.013

11. Bozkurt TO, Schornack S, Banfield MJ, Kamoun S. Oomycetes, effectors, and all that jazz. hCurr Opin Plant Biol. Elsevier Ltd; 2012;15: 483–92. doi:10.1016/j.pbi.2012.03.008

12. Jones JDG, Dangl JL. The plant immune system. Nature. 2006;444: 323–329. doi:10.1038/nature05286

13. Birch PRJ, Armstrong M, Bos J, Boevink P, Gilroy EM, Taylor RM, et al. Towards understanding the virulence functions of RXLR effectors of the oomycete plant pathogen Phytophthora infestans. J Exp Bot. 2009;60: 1133–1140. doi:10.1093/jxb/ern353

14. Whisson SC, Boevink PC, Moleleki L, Avrova AO, Morales JG, Gilroy EM, et al. A translocation signal for delivery of oomycete effector proteins into host plant cells. Nature. 2007;450: 115–118. doi:10.1038/nature06203

15. Wawra S, Trusch F, Matena A, Apostolakis K, Linne U, Zhukov I, et al. The RxLR Motif of the Host Targeting Effector AVR3a of Phytophthora infestans Is Cleaved Before Secretion. Plant Cell. 2017;29: tpc.00552.2016. doi:10.1105/tpc.16.00552

16. Hiller NL, Hiller NL, Bhattacharjee S, Ooij C Van, Liolios K, Harrison T, et al. A Host-Targeting Signal in Virulence Proteins Reveals a Secretome in Malarial Infection. Science (80-). 2012;1934: 1934–1938. doi:10.1126/science.1102737

17. Coffey MJ, Sleebs BE, Uboldi AD, Garnham A, Franco M, Marino ND, et al. An aspartyl protease defines a novel pathway for export of Toxoplasma proteins into the host cell. Elife. 2015;4: 1–34. doi:10.7554/eLife.10809

18. Boddey JA, O’Neill MT, Lopaticki S, Carvalho TG, Hodder AN, Nebl T, et al. Export of malaria proteins requires co-translational processing of the PEXEL motif independent of phosphatidylinositol-3-phosphate binding. Nat Commun. 2016;7. doi:10.1038/ncomms10470

19. Thines M. Bridging the gulf: Phytophthora and downy mildews are connected by rare grass parasites. Ausubel FM, editor. PLoS One. Public Library of Science; 2009;4: e4790. doi:10.1371/journal.pone.0004790

20. Bailey K, Cevik V, Holton N, Byrne-Richardson J, Sohn KH, Coates M, et al. Molecular cloning of ATR5(Emoy2) from Hyaloperonospora arabidopsidis, an avirulence determinant that triggers RPP5-mediated defense in Arabidopsis. Mol Plant Microbe Interact. 2011;24: 827–838. doi:10.1094/MPMI-12-10-0278

21. Tian M, Win J, Savory E, Burkhardt A, Held M, Brandizzi F, et al. 454 Genome sequencing of Pseudoperonospora cubensis reveals effector proteins with a QXLR translocation motif. Mol Plant Microbe Interact. The American Phytopathological Society; 2011;24: 543–53. doi:10.1094/MPMI-08-10-0185

22. Stassen J, Seidl M. Effector identification in the lettuce downy mildew Bremia lactucae by massively parallel transcriptome sequencing. Mol Plant Pathol. 2012;13: 719–31. doi:10.1111/j.1364-3703.2011.00780.x

23. Stassen JHM, Boer E den, Vergeer PWJ, Andel A, Ellendorff U, Pelgrom K, et al. Specific In Planta Recognition of Two GKLR Proteins of the Downy Mildew Bremia lactucae Revealed in a Large Effector Screen in Lettuce. Mol Plant Microbe Interact. The American Phytopathological Society; 2013;26: 1259–1270. doi:10.1094/MPMI-05-13-0142-R

24. Sharma R, Xia X, Cano LM, Evangelisti E, Kemen E, Judelson H, et al. Genome analyses of the sunflower pathogen Plasmopara halstedii provide insights into effector evolution in downy mildews and Phytophthora. BMC Genomics. BMC Genomics; 2015;16: 741. doi:10.1186/s12864-015-1904-7

25. Jiang RHY, Tripathy S, Govers F, Tyler BM. RXLR effector reservoir in two Phytophthora species is dominated by a single rapidly evolving superfamily with more than 700 members. Proc Natl Acad Sci U S A. 2008;105: 4874–4879. doi:10.1073/pnas.0709303105

26. Win J, Krasileva K V, Kamoun S, Shirasu K, Staskawicz BJ, Banfield MJ. Sequence divergent RXLR effectors share a structural fold conserved across plant pathogenic oomycete species. PLoS Pathog. 2012;8: e1002400. doi:10.1371/journal.ppat.1002400

27. Boutemy L, King S, Win J, Hughes R, Clarke T, Blumenschein T, et al. Structures of Phytophthora RXLR effector proteins a conserved but adaptable fold underpins functional diversity. J Biol Chem. 2011;286: 35834–42. doi:10.1074/jbc.M111.262303

28. Chou S, Krasileva K V, Holton JM, Steinbrenner AD, Alber T, Staskawicz BJ. Hyaloperonospora arabidopsidis ATR1 effector is a repeat protein with distributed recognition surfaces. Proc Natl Acad Sci U S A. 2011;108: 13323–13328. doi:10.1073/pnas.1109791108

29. Yaeno T, Li H, Chaparro-Garcia A, Schornack S, Koshiba S, Watanabe S, et al. Phosphatidylinositol monophosphate-binding interface in the oomycete RXLR effector AVR3a is required for its stability in host cells to modulate plant immunity. Proc Natl Acad Sci. National Academy of Sciences; 2011;108: 14682–14687. doi:10.1073/PNAS.1106002108

30. He J, Ye W, Choi DS, Wu B, Zhai Y, Guo B, et al. Structural analysis of *Phytophthora* suppressor of RNA silencing 2 (PSR2) reveals a conserved modular fold contributing to virulence. Proc Natl Acad Sci. 2019;2: 201819481. doi:10.1073/pnas.1819481116

31. Dou D, Kale SD, Wang X, Chen Y, Wang Q, Wang X, et al. Conserved C-terminal motifs required for avirulence and suppression of cell death by Phytophthora sojae effector Avr1b. Plant Cell. 2008;20: 1118–1133. doi:10.1105/tpc.107.057067

32. Bos JIB, Armstrong MR, Gilroy EM, Boevink PC, Hein I, Taylor RM, et al. Phytophthora infestans effector AVR3a is essential for virulence and manipulates plant immunity by stabilizing host E3 ligase CMPG1. Proc Natl Acad Sci U S A. National Academy of Sciences; 2010;107: 9909–14. doi:10.1073/pnas.0914408107

33. King SRF, McLellan H, Boevink PC, Armstrong MR, Bukharova T, Sukarta O, et al. Phytophthora infestans RXLR Effector PexRD2 Interacts with Host MAPKKK{varepsilon} to Suppress Plant Immune Signaling. Plant Cell. 2014;26: 1345–59. doi:10.1105/tpc.113.120055

34. Maqbool A, Hughes RK, Dagdas YF, Tregidgo N, Zess E, Belhaj K, et al. Structural basis of host Autophagy-related protein 8 (ATG8) binding by the Irish potato famine pathogen effector protein PexRD54. J Biol Chem. 2016;8: jbc.M116.744995. doi:10.1074/jbc.M116.744995

35. Guo B, Wang H, Yang B, Jiang W, Jing M, Li H, et al. Phytophthora sojae effector PsAvh240 inhibits a host aspartic protease secretion to promote infection. Mol Plant. Chinese Society for Plant Biology; 2019; doi:10.1016/j.molp.2019.01.017

36. Qiao Y, Liu L, Xiong Q, Flores C, Wong J, Shi J, et al. Oomycete pathogens encode RNA silencing suppressors. Nat Genet. Nature Publishing Group; 2013;45: 330–3. doi:10.1038/ng.2525

37. Xiong Q, Ye W, Choi D, Wong J, Qiao Y, Tao K, et al. Phytophthora Suppressor of RNA Silencing 2 Is a Conserved RxLR Effector that Promotes Infection in Soybean and Arabidopsis thaliana. Mol Plant- …. 2014;27: 1379–1389. Available: http://apsjournals.apsnet.org/doi/abs/10.1094/MPMI-06-14-0190-R

38. Hou Y, Zhai Y, Feng L, Karimi HZ, Rutter BD, Zeng L, et al. A Phytophthora Effector Suppresses Trans-Kingdom RNAi to Promote Disease Susceptibility. Cell Host Microbe. Elsevier Inc.; 2019;25: 153–165.e5. doi:10.1016/j.chom.2018.11.007

39. Goritschnig S, Steinbrenner AD, Grunwald DJ, Staskawicz BJ. Structurally distinct Arabidopsis thaliana NLR immune receptors recognize tandem WY domains of an oomycete effector. New Phytol. 2016;210: 984–96.

40. Fletcher K, Gil J, Bertier LD, Kenefick A, Wood KJ, Zhang L, et al. Genomic signatures of heterokaryosis in the oomycete pathogen Bremia lactucae. Nat Commun. 2019;10: 2645. doi:10.1101/516526

41. Pelgrom AJE, Eikelhof J, Elberse J, Meisrimler CN, Raedts R, Klein J, et al. Recognition of lettuce downy mildew effector BLR38 in Lactuca serriola LS102 requires two unlinked loci. Molecular Plant Pathology. 2018: 240–253. doi:10.1111/mpp.12751

42. Shen D, Li Q, Sun P, Zhang M, Dou D. Intrinsic disorder is a common structural characteristic of RxLR effectors in oomycete pathogens. Fungal Biol. Elsevier Ltd; 2017;121: 911–919. doi:10.1016/j.funbio.2017.07.005

43. Maisonneuve B. Lactuca virosa, a source of disease resistance genes for lettuce breeding: results and difficulties for gene introgression. Eucarpia leafy Veg ‘03. 2003;2003: 61–67. Available:http://www.leafyvegetables.nl/download/12_061-067_maisonneuve.pdf

44. Brameier M, Krings A, MacCallum RM. NucPred - Predicting nuclear localization of proteins. Bioinformatics. 2007;23: 1159–1160. doi:10.1093/bioinformatics/btm066

45. Anderson RG, Deb D, Fedkenheuer K, Mcdowell JM. Recent Progress in RXLR Effector Research. Mol Plant Microbe Interact. 2015;28: 1063–1072. doi:10.1094/MPMI-01-15-0022-CR

46. Marin M, Uversky VN, Ott T. Intrinsic Disorder in Pathogen Effectors: Protein Flexibility as an Evolutionary Hallmark in a Molecular Arms Race. Plant Cell. 2013;25: 3153–3157. doi:10.1105/tpc.113.116319

47. Uversky VN. Intrinsic disorder-based protein interactions and their modulators. Curr Pharm Des. 2013;19: 4191–213. Available: http://www.ncbi.nlm.nih.gov/pubmed/23170892

48. Iakoucheva LM, Brown CJ, Lawson JD, Obradović Z, Dunker AK. Intrinsic Disorder in Cell-signaling and Cancer-associated Proteins. J Mol Biol. 2002;323: 573–584. doi:10.1016/S0022-2836(02)00969-5

49. Meyers BC, Kozik A, Griego A, Kuang H, Michelmore RW. Genome-Wide Analysis of NBS-LRR – Encoding Genes in Arabidopsis. 2003;15: 809–834. doi:10.1105/tpc.009308.During

50. Hofberger JA, Jones JDG. A Novel Approach for Multi-Domain and Multi-Gene Family Identification Provides Insights into Evolutionary Dynamics of Disease Resistance Genes in Core Eudicot Plants Netherlands Chinese Academy of Sciences / Max Planck Partner Institute for Computational. 2014;31: 1–35.

51. He Q, McLellan H, Hughes RK, Boevink PC, Armstrong M, Lu Y, et al. *Phytophthora infestans* effector SFI 3 targets potato UBK to suppress early immune transcriptional responses. New Phytol. John Wiley & Sons, Ltd (10.1111); 2018; nph.15635. doi:10.1111/nph.15635

52. Vleeshouwers VGAA, Oliver RP. Effectors as tools in disease resistance breeding against biotrophic, hemibiotrophic, and necrotrophic plant pathogens. Mol Plant Microbe Interact. 2014;27: 196–206. doi:10.1094/MPMI-10-13-0313-IA

53. Giesbers AKJ, Pelgrom AJE, Visser RGF, Niks RE, Van den Ackerveken G, Jeuken MJW. Effector-mediated discovery of a novel resistance gene against *Bremia lactucae* in a nonhost lettuce species. New Phytol. 2017; doi:10.1111/nph.14741

54. Giesbers AKJ, Boer E Den, Braspenning DNJ, Bouten TPH, Specken JW, van Kaauwen MPW, et al. Bidirectional backcrosses between wild and cultivated lettuce identify loci involved in nonhost resistance to downy mildew. Theor Appl Genet. Springer Berlin Heidelberg; 2018; 1–16. doi:10.1007/s00122-018-3112-8

55. Leonelli L, Pelton J, Schoeffler A, Dahlbeck D, Berger J, Wemmer DE, et al. Structural elucidation and functional characterization of the Hyaloperonospora arabidopsidis effector protein ATR13. Tyler B, editor. PLoS Pathog. Public Library of Science; 2011;7: e1002428. doi:10.1371/journal.ppat.1002428

56. Michelmore RW, Christopoulou M, Caldwell KS. Impacts of resistance gene genetics, function, and evolution on a durable future. Annu Rev Phytopathol. 2013;51: 291–319. doi:10.1146/annurev-phyto-082712-102334

57. Eddy SR. Accelerated Profile HMM Searches. PLoS Computational Biology. 2011. p. e1002195. doi:10.1371/journal.pcbi.1002195

58. Petersen TN, Brunak S, von Heijne G, Nielsen H. SignalP 4.0: discriminating signal peptides from transmembrane regions. Nat Methods. Nature Publishing Group; 2011;8: 785–786. doi:10.1038/nmeth.1701

59. Käll L, Krogh A, Sonnhammer ELL. Advantages of combined transmembrane topology and signal peptide prediction-the Phobius web server. Nucleic Acids Res. 2007;35: 429–432. doi:10.1093/nar/gkm256

60. Peng K, Radivojac P, Vucetic S, Dunker AK, Obradovic Z. Length-dependent prediction of protein in intrinsic disorder. BMC Bioinformatics. 2006;7: 1–17. doi:10.1186/1471-2105-7-208

61. Edgar RC. MUSCLE: multiple sequence alignment with high accuracy and high throughput. Nucleic Acids Res. 2004;32: 1792–7. doi:10.1093/nar/gkh340

62. Letunic I, Bork P. Interactive Tree Of Life (iTOL): an online tool for phylogenetic tree display and annotation. Bioinformatics. 2007;23: 127–8. doi:10.1093/bioinformatics/btl529

63. Altschul SF, Gish W, Miller W, Myers EW, Lipman DJ. Basic local alignment search tool. J Mol Biol. 1990;215: 403–410. doi:10.1016/S0022-2836(05)80360-2

64. Baxter L, Tripathy S, Ishaque N, Boot N, Cabral A, Kemen E, et al. Signatures of adaptation to obligate biotrophy in the Hyaloperonospora arabidopsidis genome. Science (80-). 2010;330: 1549–1551. doi:10.1126/science.1195203

65. Lamour KH, Mudge J, Gobena D, Hurtado-Gonzales OP, Schmutz J, Kuo A, et al. Genome sequencing and mapping reveal loss of heterozygosity as a mechanism for rapid adaptation in the vegetable pathogen Phytophthora capsici. [Internet]. Molecular plant-microbe interactions: MPMI. 2012. pp. 1350–60. doi:10.1094/MPMI-02-12-0028-R

66. Broad Institute. Phytophthora parasitica INRA-310 Genome sequencing and assembly. [Internet]. 2018 [cited 6 Jun 2018]. Available: https://data.nal.usda.gov/dataset/phytophthora-parasitica-inra-310-genome-sequencing-and-assembly

67. Jiang RHY, de Bruijn I, Haas BJ, Belmonte R, Löbach L, Christie J, et al. Distinctive Expansion of Potential Virulence Genes in the Genome of the Oomycete Fish Pathogen Saprolegnia parasitica. PLoS Genet. 2013;9: e1003272. doi:10.1371/journal.pgen.1003272

68. Yin L, An Y, Qu J, Li X, Zhang Y, Dry I, et al. Genome sequence of Plasmopara viticola and insight into the pathogenic mechanism. Sci Rep. Nature Publishing Group; 2017;7: 1–12. doi:10.1038/srep46553

69. Dussert Y, Couture C, Mazet ID, Gouzy J, Piron M-C, Kuchly C, et al. Genome assembly and annotation of Plasmopara viticola, the grapevine downy mildew pathogen [Internet]. Portail Data Inra; 2019. doi:doi/10.15454/4NYHD6

70. Derevnina L, Reyes-Chin Wo S, Martin F, Wood K, Froenicke L, Spring O, et al. Genome Sequence and Architecture of the Tobacco Downy Mildew Pathogen, Peronospora tabacina. Mol plant-microbe Interact. 2015;28: 1198–1215. doi:10.1094/MPMI-05-15-0112-R

71. Fletcher K, Klosterman SJ, Derevnina L, Martin F, Bertier LD, Koike S, et al. Comparative genomics of downy mildews reveals potential adaptations to biotrophy. BMC Genomics. BMC Genomics; 2018;19: 8–10. doi:10.1186/s12864-018-5214-8

72. Zhong S, Joung JG, Zheng Y, Chen YR, Liu B, Shao Y, et al. High-throughput illumina strand-specific RNA sequencing library preparation. Cold Spring Harb Protoc. 2011;6: 940–949. doi:10.1101/pdb.prot5652

73. Andrews S. FastQC: a quality control tool for high throughput sequence data [Internet]. 2010.

74. Li H. Aligning sequence reads, clone sequences and assembly contigs with BWA-MEM. 2013;00: 1–3. doi:10.1186/s13756-018-0352-y

75. Dobin A, Davis CA, Schlesinger F, Drenkow J, Zaleski C, Jha S, et al. STAR: Ultrafast universal RNA-seq aligner. Bioinformatics. 2013;29: 15–21. doi:10.1093/bioinformatics/bts635

76. Warnes GR, Bolker B, Bonebakker L, Gentleman R, Huber W, Liaw A, et al. Package ‘gplots’ [Internet]. 2012. Available: http://cran.r-project.org

77. Team RC. R: A language and environment for statistical computing. [Internet]. Vienna, Austria.: R Foundation for Statistical Computing; 2014. Available: http://www.r-project.org/

78. Earley KW, Haag JR, Pontes O, Opper K, Juehne T, Song K, et al. Gateway-compatible vectors for plant functional genomics and proteomics. Plant J. 2006;45: 616–629. doi:10.1111/j.1365-313X.2005.02617.x

79. Wroblewski T, Tomczak A, Michelmore R. Optimization of Agrobacterium-mediated transient assays of gene expression in lettuce, tomato and Arabidopsis. Plant Biotechnol J. 2005;3: 259–273. Available: http://www.ncbi.nlm.nih.gov/pubmed/17173625

80. Wroblewski T, Caldwell KS, Piskurewicz U, Cavanaugh KA, Xu H, Kozik A, et al. Comparative large-scale analysis of interactions between several crop species and the effector repertoires from multiple pathovars of Pseudomonas and Ralstonia. Plant Physiol. 2009;150: 1733–49. doi:10.1104/pp.109.140251

81. Sperschneider J, Catanzariti AM, Deboer K, Petre B, Gardiner DM, Singh KB, et al. LOCALIZER: Subcellular localization prediction of both plant and effector proteins in the plant cell. Sci Rep. Nature Publishing Group; 2017;7: 1–14. doi:10.1038/srep44598

82. Felix G, Duran JD, Volko S, Boller T. Plants have a sensitive perception system for the most conserved domain of bacterial flagellin. Plant J. 1999;18: 265–276. doi:10.1046/j.1365-313X.1999.00265.x

83. Jamir Y, Guo M, Oh HS, Petnicki-Ocwieja T, Chen S, Tang X, et al. Identification of Pseudomonas syringae type III effectors that can suppress programmed cell death in plants and yeast. Plant J. 2004;37: 554–565. doi:10.1046/j.1365-313X.2003.01982.x

