## Supplemental Figures and Tables for "Effector prediction and characterization in the oomycete pathogen *Bremia lactucae* reveal host-recognized WY domain proteins that lack the canonical RXLR motif"

Supplemental Information

**Supplemental Table 1.** RXLR-like sequences observed within 100 amino acids of the N-terminus of *B. lactucae* WY proteins, sorted by first amino acid. Sequences that occur in more than one protein are highlighted in red.

| *Starting with:* | **Glycine** | **Histidine** | **Lysine** | **Glutamine** | **Arginine** |
|---|---|---|---|---|---|
| | GCLR | HAVN | KALN | QAIN | RAVN |
| | GDMN | HFR | KDFK | QEIN | RFK |
| | GELK | HIK | KDLN | QELR | RFN |
| | GHLK | HILN | KFK | QFN | RFR |
| | GKMK | HIR | KFMN | QGVR | RFVK |
| | GLFK | HLK | KIK | QIIK | RHLN |
| | GLK | HLN | KIMR | QLN | RIR |
| | GLLN | HLR | KIR | QLR | RIYR |
| | GLN | HMN | KIVK | QLVK | RIYR |
| | GVK | HSFK | KKLK | QLVN | RLK |
| | | HSVN | KLIK | QSIR | RLR |
| | | HYVK | KLK | QTFK | RMLK |
| | | | KLN | QTFN | RMN |
| | | | KLR | QVVR | RPLK |
| | | | KMFK | QVYK | RPVN |
| | | | KRLK | QYLK | RQFN |
| | | | KSFK | | RRLN |
| | | | KSVK | | RSIR |
| | | | KSVN | | RSLK |
| | | | KVFK | | RTIK |
| | | | KVK | | RVFK |
| | | | KVLN | | RVK |
| | | | KVN | | RVR |
| | | | KVR | | RYLK |
| | | | KYR | | RYNK |

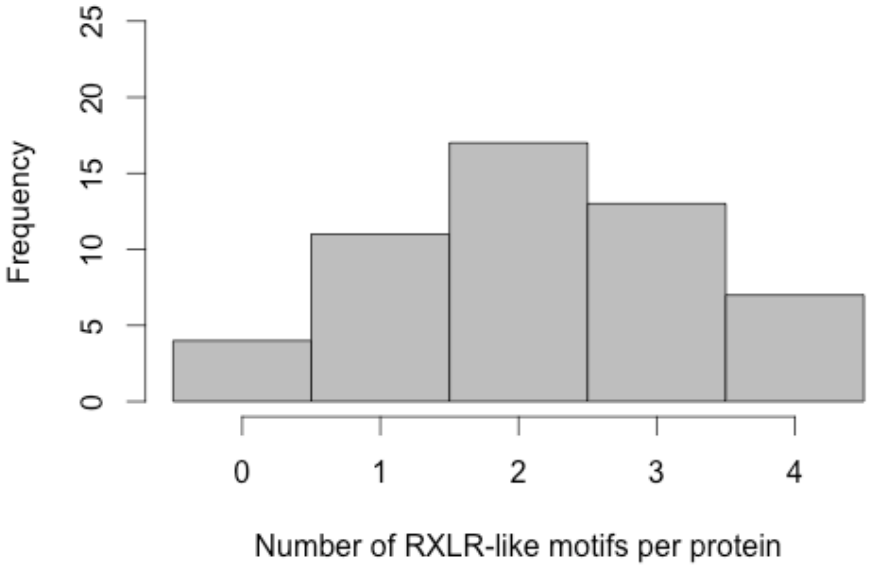


**Supplemental Figure 1.** The number of RXLR-like motifs detected in the first 100 amino acids of the N-terminus of *B. lactucae* WY effectors.

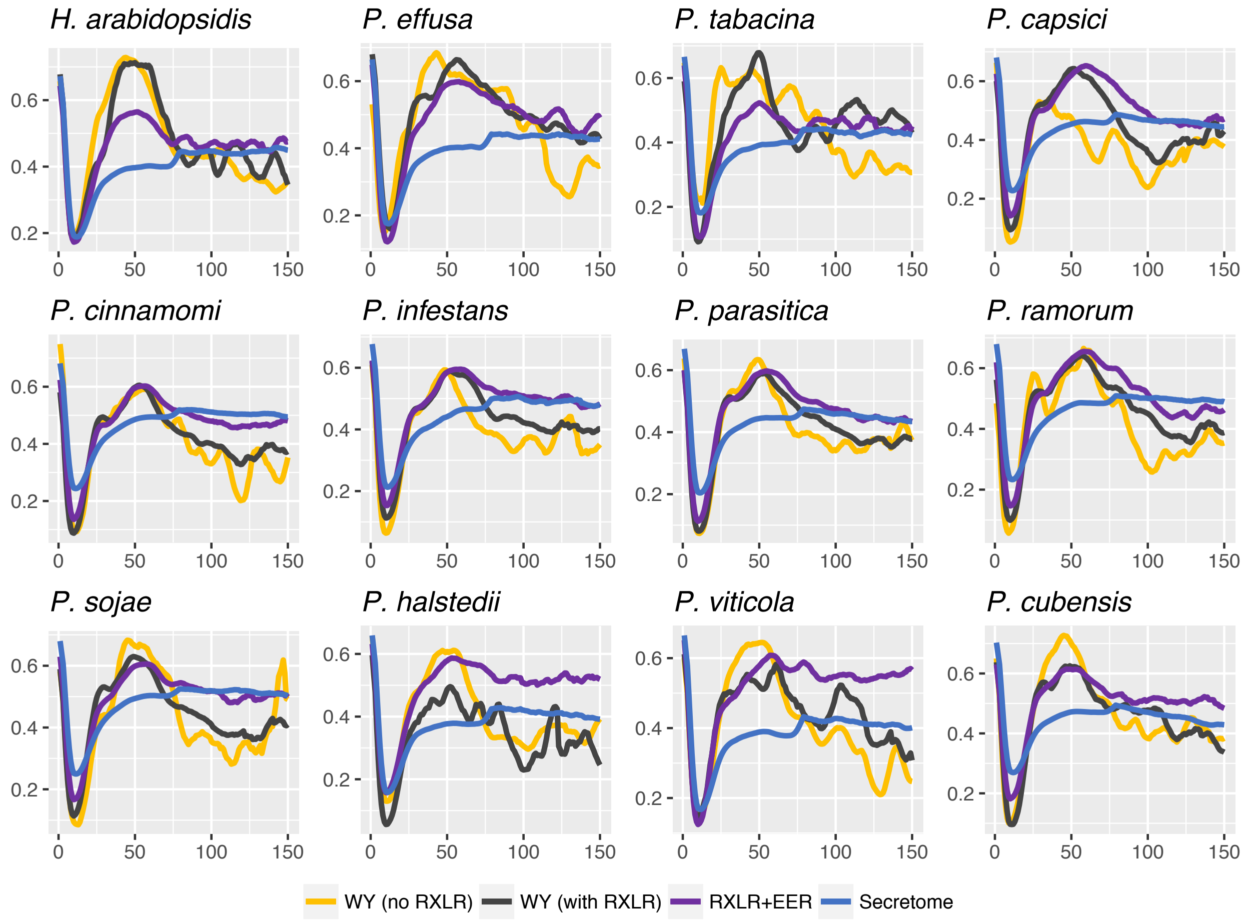


**Supplemental Figure 2.** Intrinsic disorder of first 150 amino acids of RXLR and WY containing candidate effectors mined from other oomycete genomes. The x-axis shows the amino acid position; the y-axis shows the intrinsic disorder (VSL2 PONDR) score. Proteins were categorized as WY with no RXLR (yellow), WYs with RXLR (black), RXLR+EER (with or without WY, purple), and the total predicted secretome (blue) using the same color scheme as Fig. 2. RXLR and EER motifs were as described in Figs. 2 and 3. Average positional disorder scores are shown for each class of proteins.

**A**

| Genotype | Species | BLN06 | GFP | +control |
|---|---|---|---|---|
| CGN14312 | indica | 3.3 | 1.5 | 3.8 |
| LS102 | serriola | 0.0 | 0.0 | 4.0 |
| NunDm17 | sativa | 0.8 | 0.0 | 4.0 |
| RYZ2164 | sativa | 0.0 | 0.0 | |

**B**

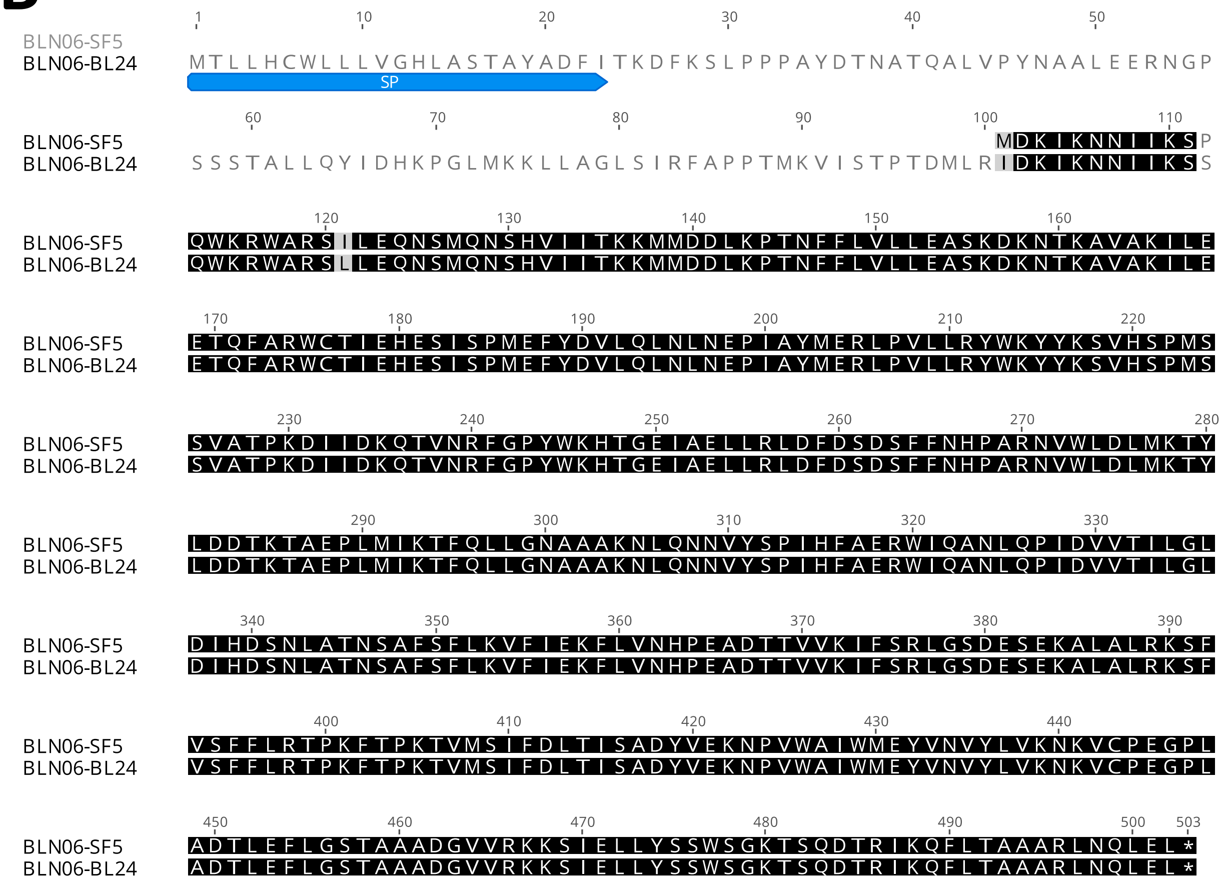


**Supplemental Figure 3.** Agroinfiltration results from BLN06-SF5 (A) and sequence comparison between BLN06 between SF5 and BL24 (B) [1].


1. Pelgrom AJE, Eikelhof J, Elberse J, Meisrimler CN, Raedts R, Klein J, et al. Recognition of lettuce downy mildew effector BLR38 in Lactuca serriola LS102 requires two unlinked loci. Molecular Plant Pathology. 2018: 240–253. doi:10.1111/mpp.12751
